# The Functional Connectome Mediating Circadian Synchrony in the Suprachiasmatic Nucleus

**DOI:** 10.1101/2024.12.06.627294

**Authors:** K.L. Nikhil, Bharat Singhal, Daniel Granados-Fuentes, Jr-Shin Li, István Z. Kiss, Erik D. Herzog

## Abstract

Circadian rhythms in mammals arise from the spatiotemporal synchronization of ∼20,000 neuronal clocks in the Suprachiasmatic Nucleus (SCN). While anatomical, molecular, and genetic approaches have revealed diverse cell types and signaling mechanisms, the network wiring that enables SCN cells to communicate and synchronize remains unclear. To overcome the challenges of revealing functional connectivity from fixed tissue, we developed MITE (Mutual Information & Transfer Entropy), an information theory approach that infers directed cell-cell connections with high fidelity. By analyzing 3447 hours of continuously recorded clock gene expression from 9011 cells in 17 mice, we found that the functional connectome of SCN was highly conserved bilaterally and across mice, sparse, and organized into a dorsomedial and a ventrolateral module. While most connections were local, we discovered long-range connections from ventral cells to cells in both the ventral and dorsal SCN. Based on their functional connectivity, SCN cells can be characterized as circadian signal generators, broadcasters, sinks, or bridges. For example, a subset of VIP neurons acts as hubs that generate circadian signals critical to synchronize daily rhythms across the SCN neural network. Simulations of the experimentally inferred SCN networks recapitulated the stereotypical dorsal-to-ventral wave of daily PER2 expression and ability to spontaneously synchronize, revealing that SCN emergent dynamics are sculpted by cell-cell connectivity. We conclude that MITE provides a powerful method to infer functional connectomes, and that the conserved architecture of cell-cell connections mediates circadian synchrony across space and time in the mammalian SCN.

**Highlights:**

- We developed MITE, an information theory method, to accurately infer directed functional connectivity among circadian cells.
- SCN cell types with conserved connectivity patterns spatially organize into two regions and function as generators, broadcasters, sinks, or bridges of circadian information.
- One-third of VIP neurons serve as hubs that drive circadian synchrony across the SCN.
- Key connectivity features mediate the generation and maintenance of intercellular synchrony and daily waves of clock gene expression across the SCN.

## Introduction

Circadian clocks are biological mechanisms that drive daily changes—called circadian rhythms—in vital processes such as sleep, metabolism, immune responses, and cell division, in alignment with solar day-night cycles^1–7^. In mammals, ∼20,000 neuronal clocks in the hypothalamic Suprachiasmatic Nucleus (SCN) drive these daily rhythms^8^. These neuronal clocks intrinsically generate autonomous rhythmic gene expression and firing patterns through a conserved transcriptional-translational feedback loop that operates with ∼24-hour periodicity^8–10^. However, individual SCN neurons are sloppy cellular clocks^11^ so that, when cells can’t communicate, they lose coherence and amplitude of their circadian rhythms^12–15^. When SCN cells synchronize, they exhibit a high amplitude, precise, wave of circadian gene expression that starts in the dorsal region and moves ventrally to drive daily rhythms in physiology and behavior^12–16^. Revealing the anatomical and functional neural interactions that underlie such emergent behavior remains a major challenge in neuroscience^17–20^. This study tests for cell-cell connectivity features that generate and sustain synchronized circadian rhythms in the SCN.

Based on gene and protein expression profiling, the SCN has been described to comprise 8 cell classes and 11 transcriptionally distinct cell types expressing over 25 neuropeptides organized into a dorsal ’shell’, dominated by vasopressin (AVP) neurons, and a ventral ’core’, which primarily contains vasoactive intestinal peptide (VIP) and gastrin-releasing peptide (GRP) expressing neurons^8,21–31^. Tracing anatomical projections and light responses in SCN cells has led to a canonical model in which the retinorecipient core region signals photic information to the shell, synchronizing the organism to day-night cycles^23,32–38^. Pharmacology and genetic manipulations have placed VIP as a critical synchronizer of SCN cells^29,39–47^. However, other neuropeptides (e.g., GRP and Prokineticin, PROK2) serve as weaker synchronizers, as evidenced by a subset of mice that retain circadian behaviors even after loss of SCN VIP, its receptor (VPAC2) or the neurons^16,23,25,26,29,30,48^. Furthermore, contrary to their previously believed roles as circadian output cells, the AVP neurons were recently found to exhibit pacemaker-like properties and couple SCN rhythms^48–55^. In one striking example, SCN explants did not synchronize when molecular clocks were disrupted within AVP, but not the VIP neurons^52^, suggesting that circadian synchrony depends on SCN AVP cells. Here, we aimed to test whether connectivity patterns are cell-type specific and important for SCN timekeeping.

Electron microscopy and fluorescence imaging have aided the construction of anatomical complete connectomes and testable predictions about neuronal function in smaller organisms like *Caenorhabditis*^56^ and *Drosophila*^57,58^, and about 0.2% of the mouse brain^59–63^. With the goal of complementing structural maps of intercellular communication, there have been efforts to map cell-cell functional connectomes^17–19^ based on physiological data^64–68^. For example, one study used circadian gene expression of synchronizing SCN cells and identified sparse, undirected connections which are higher in and between the bilateral core regions^67^. Another study found that GABA-dependent firing patterns revealed a distinct, faster cell-cell communication network that changes strength with time of day and weakly opposes synchronization of circadian rhythms^68^. Modelling studies have predicted a wide range of possible SCN connectivity topologies including, small-world^69^, mean-field^70,71^, nearest-neighbor^72^ and mix of multiple topologies^73^, each shaping SCN dynamics. An essential next step towards understanding how network topology mediates these dynamics requires mapping cell-cell connectivity to cell types and their functions. Because data limitations hindered accurate inference of directed connections using existing methods^64,74,75^, we hypothesized that novel methods and data sets would allow us to reveal the directed cell-cell connectivity that mediates daily rhythms in the SCN.

## Results

### MITE Infers SCN Cell-Cell Connectivity with High Fidelity

Based on the premise that connected cells share causally related clock-gene dynamics during synchronization of their circadian rhythms, we developed a method to map SCN cell-cell connectivity from real-time recordings of PERIOD2 (PER2) expression. We compared three undirected (Dynamic Time Warping, DTW^76^; Maximal Information Coefficient, MIC^77^; and Mutual Information, MI^78^) and directed methods (Algorithm for Revealing Network Interactions, ARNI^79^; Granger Causality, GC^80^; and Transfer Entropy, TE^81^), each with their assumptions about causality and propensity to capture non-linear interactions among cells.

We first tested the ability of these methods to infer true connectivity from 300 simulations of 30 large circadian networks (600 cells/network, 10 simulations/network) with small-world, scale-free, or mixed topologies. Each simulated cell comprised a transcription-based model of a circadian oscillator^82^ with hourly PER2 expression recorded over 6 days to emulate experimental conditions (Supplementary Fig. 1a-c). Area Under the Receiver Operator Characteristic (AUROC), comparing the inferred connectivity with true networks revealed that, among undirected methods, MI outperformed the others by up to 37% (had higher AUROC; Supplementary Figs. 1d and 2) and required at least five times less computation time, while TE outperformed others by up to 39.4% among the directed methods (Supplementary Figs. 1d and 3). Reasoning that high MI and TE values together reflect most likely connections between cells, we multiplied the two values for each pairwise cell-cell interaction to generate MITE scores (Mutual Information & Transfer Entropy, See Methods). We found that MITE consistently outperformed TE with up to 39.8% higher AUROC across multiple network topologies (Supplementary Figs. 1e and 3).

Because SCN cells can spontaneously recover circadian synchrony after TTX-induced decoupling in vitro^15,67,68^, we next tested MITE on movies of cellular PER2 expression in synchronizing SCN explants recorded before, during, and after TTX treatment. Consistent with prior observations^15,67,68^, TTX addition induced SCN cells to lose coherent daily rhythms in PER2 expression which was gradually regained over several days after TTX removal (Fig. 1a). Similar to a strategy employed earlier^67,68^, we used PER2 traces of synchronizing SCN cells and based on comparisons of 9.49 million connections across 3081 cells, we computed the z-transformed MITE scores for all possible cellular interactions, removed connections with z-scores below the empirically computed threshold for interactions between cells in separate SCN (i.e., impossible connections), and revealed SCN networks with low false discovery rates (below 0.05, defined as probability of falsely identifying an impossible SCN connection) and high hit-rates (above 95%, defined as percent of likely true intra-SCN connections among all identified connections; Fig. 1b-d and Supplementary Fig. 4a,b).

**Fig. 1.**
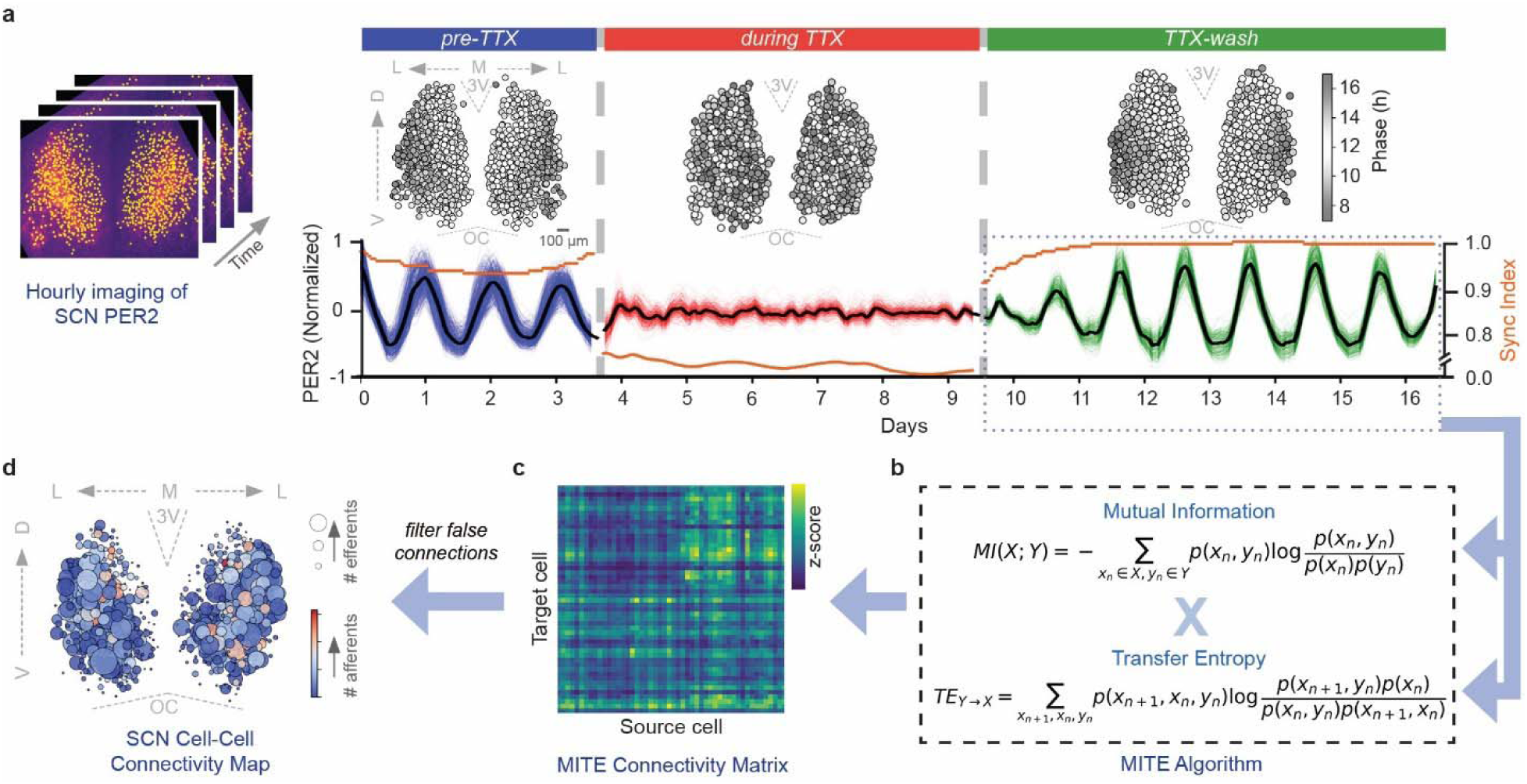
Pipeline for inferring functional connectivity among cells of the Suprachiasmatic Nucleus (SCN). **a**, Longitudinal hourly bioluminescence imaging of cells (yellow) within a representative PERIOD2::LUCIFERASE SCN explant (left) generates PER2 expression (right) before, during, and after tetrodotoxin (TTX) treatment. PER2 expression (black line=mean) and synchrony index (orange line) calculated from the hundreds of SCN cells (blue, red, and green lines) illustrate how circadian cells spontaneously resynchronize after TTX removal. Phase maps show how the dorsal-to-ventral wave of daily cellular PER2 expression was abolished by TTX and restored after TTX removal. **b,** Using our Mutual Information & Transfer Entropy (MITE) algorithm, we processed the PER2 traces recorded during resynchronization (dotted blue box) to infer MITE connectivity matrix **(c)** of stronger (yellow) and weaker (blue) pairwise cellular interactions. **d**, After thresholding the connectivity matrix based on the empirically computed z-score of known false connections, we revealed the functional cell-cell connectivity map of SCN with more efferents (larger circles) or afferents (warmer colors). D, Dorsal; V, Ventral; M, Medial; L, Lateral; 3V, 3rd Ventricle; OC, Optic Chiasm.

To further assess MITE’s accuracy, we compared the bilateral SCN maps of seven mice of similar age, sex, and light-dark histories using two methods (Directed Graphlet Correlation Distance DGCD13 and DGCD129)^83,84^ and found highly conserved connectivity structure among independently recorded SCN (Supplementary Fig. 4c). These findings demonstrate that MITE accurately identifies directed cell-cell interactions that are revealed during circadian synchronization among SCN cells.

### The Bilaterally Similar SCN Network has Sparse, Efficient, and Spatially Distinct Connectivity

To understand the connectivity architecture that synchronizes SCN neural ensemble, we analyzed network features of the inferred SCN maps from seven mice. Consistent with prior findings^21,67,85–87^, SCN cells projected both ipsilaterally (within left or right SCN) and contralaterally (between left and right SCN). Ipsilateral SCN projections dominated in number (59.7 ± 1.2% vs. 40.3 ± 1.2%, ipsilateral vs. contralateral, mean ± SD, *n* = 7 SCN; Fig. 2a and Supplementary Fig. 5a and Supplementary Table 1) and strength (3% higher MITE scores for ipsilateral connections, *p* = 0.0015, Cohen’s d = 0.708). The SCN connectome was sparse, with each cell projecting to only 3.6 ± 0.3% of the network, or 719 ± 72 projections per cell (mean degree, defined as the percent of the network to which a cell connects); while some cells had high number of efferents (Fig. 2c and Supplementary Fig. 5b). The inferred networks’ structural statistics indicated efficient coupling, based on a small-worldness coefficient greater than one and relatively short path lengths of 3.4 ± 0.4 cells (average number of cells connecting any two SCN cells; Fig. 2d and Supplementary Fig. 6a and Supplementary Table 1).

**Fig. 2.**
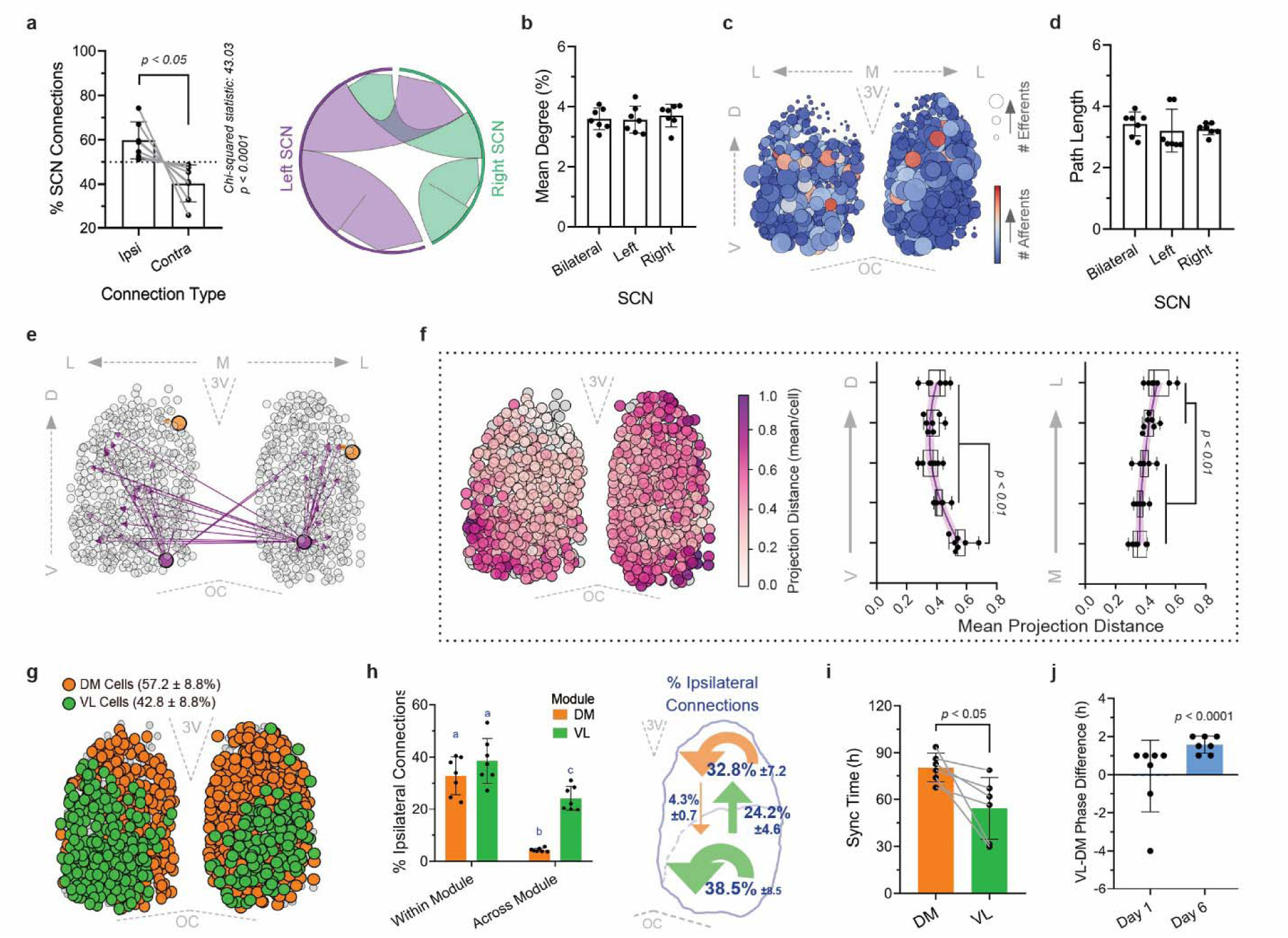
MITE reveals two networks within the SCN that differ in their connectivity and circadian properties. **a**, Cell-cell connections identified across seven SCN show more ipsilateral (about 60% of all connections were within left or right SCN) than contralateral connections between left and right SCN (paired Student’s t-test for ipsi vs. contra comparisons, *p* < 0.05; Chi-squared test for comparisons with expected frequencies of 50%, *p* < 0.0001). Ribbon plot (right) of ipsilateral (thick lavender and sea-green arrows projecting within left and right SCN) and contralateral cell-cell connections (arrows projecting across left and right SCN) identified within a representative bilateral SCN (see Supplementary Table 1 for all seven SCN). **b,** The inferred mean degree (percent of network to which a cell connects, 3.6 ± 0.3%, mean ± SD) was consistently low both in SCN of different mice and in left and right SCN of the same mouse, suggesting that the SCN connectivity is sparse and conserved (one-way ANOVA with Tukey’s post-hoc comparisons, *p* > 0.05, *n* = 7 SCN). **c**, Ventral SCN cells sent more efferents (larger circles) while most SCN cells received similar numbers of afferents (blue saturation), shown from a representative SCN (see Supplementary Fig. 5 for all seven SCN). **d,** The path length (average number of cells that connect any two cells, 3.3 ± 0.4 cells, mean ± SD) was consistently low both in SCN of different mice and in left and right SCN of the same mouse, revealing highly efficient connectivity among SCN cells (one-way ANOVA with Tukey’s post-hoc comparisons, *p* > 0.05, *n* = 7 SCN). **e,** Ventral SCN cells (purple circles) had longer and more efferents (purple arrows) compared to dorsal cells (orange circles and arrows). These four representative cells are from SCN1 (see Supplementary Figs. 5 and 7 for maps of all seven SCN). **f**, Projection distances presented as mean Euclidian distance from a cell to all its targets, in a representative SCN (left, purple saturation) and from all seven SCN presented as the mean projection distance of cells grouped in 20% bins along the ventral-dorsal and medial-lateral axes. Note that ventrolateral SCN cells sent longer functional projections reaching up to halfway across the SCN (one-way ANOVA with Tukey’s post-hoc comparisons, *n* = 7 SCN, purple lines are non-linear fits with 95% CI shaded, error bars are SD across cells in the bin). **g,** Based on their relative density of connections, SCN cells clustered into Dorsomedial (DM, orange) and Ventrolateral modules (VL, green), with only a few excluded from either (grey circles). Numbers within parentheses indicate module size (mean ± SD, percent of all cells in the module; See Supplementary Fig. 10 for clustering based on connectivity of all seven SCN). **h**, Cells projected more within their module (DM-to-DM:32.5 ± 7.5%, VL-to-VL: 38.9 ± 8.9%), but VL cells also sent six times more projections to DM cells (VL-to-DM: 24.2 ± 4.6%, DM-to-VL: 4.3 ± 0.7%; two-way ANOVA with Tukey’s post-hoc comparisons, letters indicate *p* < 0.01, *n* = 7 SCN). The schematic (right) summarizes the fraction (mean ± SD across 7 SCN) of all ipsilateral connections projecting within and across DM (orange) and VL modules (green). **i,** Circadian properties correlated with functional connections. VL cells synchronized their circadian PER2 rhythms about a day faster than DM cells (54.2 ± 19.7 h vs. 80.4 ± 9.28 h after TTX removal, VL vs. DM, paired Student’s *t*-test, *n* = 6 SCN, reliable fit was not obtained for one SCN, see Methods). **j,** After TTX removal, cells transitioned from random phase-relationships on day 1 to a reliable wave of PER2 expression with DM cells peaking 1.6 ± 0.4 h (mean ± SD) earlier than VL cells (one sample Student’s *t*-test comparing with 0 h phase difference, *n* = 6 SCN). V, Ventral; M, Medial; L, Lateral; 3V, 3rd Ventricle; OC, Optic Chiasm.

Because two prior studies described SCN connections as distributed either exponentially (where cells have similar number of connections)^67,88^ or as a scale-free network with some highly connected hubs^89^, we compared fits of seven distributions (power-law, exponential, Weibull, lognormal, Poisson, truncated power law and normal) using maximum likelihood estimators^90^ and found that distributions of incoming connections (number of afferents/cell) in all SCN were best fit by Weibull curves and all except one SCN had out-going connections (number of efferents/cell) best fit by Weibull curves (Supplementary Fig. 6b,c). We next evaluated long-range interactions for their potential to improve information transmission^69,91^. Intriguingly, we found that dorsal SCN cells mainly projected to nearby cells, and ventrolateral SCN cells had more and longer efferents, some signaling halfway across the SCN (Fig. 2e,f and Supplementary Fig. 5b). This pattern was consistent when we analyzed the data from each unilateral or bilateral SCN (Supplementary Fig. 7), indicating that these long-range connections were not due projections to contralateral cells. These evidence for some highly connected cells, long-range interactions and Weibull distributed connections suggest that SCN has heterogeneous numbers of connections/cell that is best described as intermediate between scale-free and exponential^92^, with a few highly connected cells.

Interestingly, comparisons using mean degree, path length and two DGCD metrics revealed that the unilateral SCN maps were highly similar to each other and also to the bilateral SCN (Fig. 2a-d and Supplementary Fig. 8). This suggests that contralateral projection patterns largely mirror ipsilateral patterns. For example, we found that ventral-to-ventral projecting SCN cells within a unilateral SCN also project to contralaterally located ventral (and not) dorsal cells. Thus, SCN cells, connected as a sparse small-world network communicate with only ∼4% of the network, typically through just 3-4 cells with spatially distinct, long-range and heterogenous connections–an architecture well suited for efficient information propagation.

### Two Asymmetrically Connected Cellular Modules Underlie SCN Synchronization

Given their spatially distinct and reproducible projection patterns, we analyzed each SCN topology using three community detection methods (Greedy Modularity^93^, Girvan-Newman^94^ and Label propagation^95^). We report communities based on Greedy Modularity because it identified communities more robustly (had lower inter-SCN variation) while also retaining information about connection directions. Since community detection groups cells based on high intra-group and low inter-group connections, we used unilateral maps to avoid artifacts caused by the less frequent contralateral projections. We consistently found three cellular communities in all SCN (Supplementary Fig. 9a). Two groups comprised the vast majority of SCN cells (Greedy Modularity: 65-88% of recorded cells, Label Propagation: 71-88%, Girvan-Newman: 72-90%, *n* = 14 unilateral SCN) and localized to the ventral or dorsal SCN (Supplementary Fig. 9b). A third, smaller group comprising about 8.8 ± 5.2% cells (mean ± SD, *n* = 14 unilateral SCN) varied in its location and was found within the dorsal group in 12 of 14 unilateral SCN. Subsequently, clustering the three communities across all SCN by their mean Out-degree (percent of the network to which a cell projects) consistently revealed two cellular modules in the ventrolateral (VL) and dorsomedial (DM) SCN (Fig. 2g and Supplementary Figs. 9c-d and 10). As a further test of modular organization of the SCN, we applied MITE to three explants that failed to resynchronize their PER2 expression after TTX removal. We did not detect the VL and DM communities in these SCN, indicating that the two-module topology is reproducible and underlies circadian synchrony in the SCN (Supplementary Fig. 11).

The VL module was smaller and more densely connected than DM module. Within each unilateral SCN, the VL module comprised 42.8 ± 10.9% of all cells and sent about 63% of all projections to cells in both the VL and DM modules (mean ± SD, *n =* 14 unilateral SCN; Fig. 2g-h). We identified a strong ventral-to-dorsal connectivity (24.2% ± 4.6% of all projections went from VL to DM cells), over six times the number of backward projections from DM cells that mostly projected locally among themselves (Fig. 2e-h and Supplementary Fig. 7). Contralateral projections followed the same pattern as well (Supplementary Fig. 10b), consistent with the bilateral similarity in their network properties (Fig. 2a-d and Supplementary Fig. 8). We thus conclude that SCN network is organized as two functionally coupled cellular modules that are hierarchically and bilaterally organized with dense projections from ventral to dorsal cells and fewer reverse connections.

### Ventral SCN Cells Synchronize Faster Than Dorsal Cells

Based on the rates at which cells synchronize and respond to light, signals from ventral SCN cells are believed to entrain the dorsal cells^35,38,96–104^, although, surprising recent findings implicated dorsal-to-ventral connectivity as critical to synchronize the SCN^48–55^. We found that VL cells stabilized their circadian period about 32% faster than DM cells after TTX removal (54.2 ± 19.2 h vs. 80.4 ± 9.6 h, VL vs. DM, mean ± SD, *n* = 6 SCN, based on Gompertz curve fits to estimate the time until mean daily period of the module varied less than 0.5-hour day-to-day; Fig. 2i and Supplementary Fig. 12). Once their circadian periods stabilized, DM cells peaked each day about 1.6 h earlier than VL cells, consistent with previously observed PER2 phase-wave (Fig. 2j and Supplementary Fig. 12a). These results suggest that, in the absence of light, ventral SCN cells synchronize to each other and then drive synchrony among the dorsal cells to produce the spatiotemporal daily dorsal-to-ventral wave of PER2 expression.

### Ventral SCN Hub Cells Generate and Broadcast Signals for Circadian Synchrony

Although over 40 cell-types have been identified in the SCN, their roles in circadian synchronization remain unknown^8,27^. Having identified the dorsomedial and ventrolateral SCN modules based on the relative density of connections within and among them, we next quantified each SCN cell’s influence based on their projection patterns to the network with six centrality measures^105–107^. The Reverse Pagerank centrality identifies "generators" of synchrony signals– cells at the top of the hierarchy that project to other influential cells, while Out-degree centrality (percent of the network to which a cell projects) highlights "broadcaster" cells with many outgoing connections that disseminate signals. We also characterized "receiver" cells with many incoming connections based on high In-degree centrality (percent of the network from which a cell receives afferents), "sink" cells where more connections converge based on Pagerank centrality, "bridge" cells that mediate information flow between distinct regions based on Betweenness centrality, and cells that either quickly transmit or receive signals based on their topological proximity (short path lengths to and from other cells in the network), measured as Closeness_out-degree_ and Closeness_in-degree_ centrality, respectively^105–107^.

We found that VL cells had higher Reverse Pagerank (0.1 ± 0.008 vs. 0.02 ± 0.007, VL vs. DM, mean ± SD, *n* = 14 unilateral SCN) and Out-degree centralities than DM cells (6.2 ± 1.3% vs. 2.6 ± 0.2%, VL vs. DM; Fig. 3a and Supplementary Fig. 13). Interestingly, these two centrality measures were uncorrelated among VL cells, indicating the presence of two, partially overlapping VL cell types (Supplementary Fig. 14). For instance, most VL cells, irrespective of where they projected to, functioned as signal generators (had high Reverse Pagerank), whereas only those that exclusively projected to both the modules were broadcasters with high Out-degree (Supplementary Fig. 13b). These VL signal generators and broadcaster cells also had high Closeness_out-degree_, indicating that signals from the ventral SCN spread through the network faster than those originating from dorsal SCN (0.6 ± 0.08 vs. 0.3 ± 0.07, VL vs. DM; Supplementary Fig. 13b). Given the evidence for highly influential connectivity and partial overlap among the generators and broadcasters, we conclude that hub cells in the VL SCN send on average over two-times more efferents than DM cells and comprise partially overlapping classes of signal generators and broadcasters

**Fig. 3.**
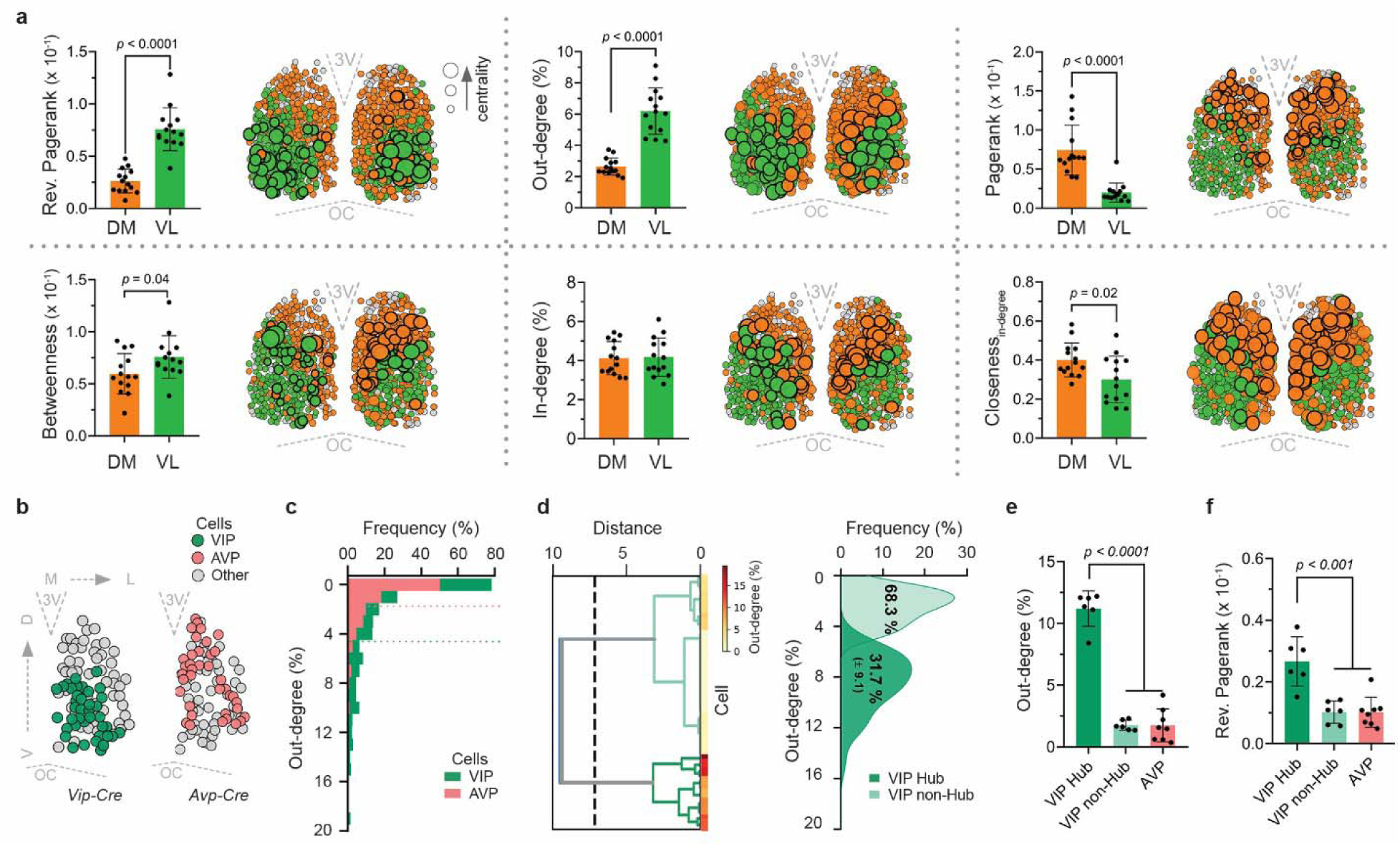
A subset of VIP neurons have highly influential connections. VL cells (green) had higher Reverse Pagerank centrality compared to DM cells (orange). This suggests VL cells act as primary circadian signal generators (top-left, larger cell = higher centrality, thick borders highlight cells in the top 30%). VL cells also had higher Out-degree centrality (suggestive of signal broadcasters, top-middle) and DM cells had higher Pagerank centrality (indicative of sink cells to which signals ultimately flow, top-right). Cells of the VL module had marginally higher Betweenness centrality (bridges mediating information flow, bottom-left). Cells of the two modules had similar In-degree (percent of the network from which a cell receives afferents, bottom-middle), whereas DM cells had higher Closeness_in-degree_ centrality (cells that are topologically closer and receive signals faster, bottom-right; unpaired Student’s t-test, *n* = 14 unilateral SCN). **b,** VIP (green) and AVP (coral) neurons subjected to MITE analysis in representative unilateral Vip-Cre (left) and Avp-Cre (right) SCN explants (grey circles = non-VIP and non-AVP cells, respectively). **c,** Frequency histograms of Out-degree (% of network to which a cell projects) for VIP (green) and AVP (coral) neurons show that VIP cells sent more projections (4.6 ± 0.2 % vs. 1.7 ± 1.1%, mean ± SD, VIP vs. AVP, *n* = 264 VIP neurons in three VIP-Cre and 572 neurons in four AVP-Cre mice). Some VIP neurons had strikingly more efferents than others (long tail in VIP histogram). **d-e,** Unsupervised hierarchical clustering revealed two groups of VIP neurons, in a representative SCN (left; See Supplementary Fig. 14 for all SCN). One group (“Hub” neurons, dark green, 31.6 ± 9.6% of all VIP neurons, mean ± SD) clustered with high Out-degree, acting as signal broadcasters by projecting to 11.2 ± 1.4% of the network, five times more than other SCN cells. The second group (“non-Hub” VIP neurons, 68.3 ± 9.6% of all VIP neurons) projected to only 1.7 ± 0.4% of SCN cells (one-way ANOVA with Tukey’s post-hoc comparisons; *n* = 6 and 8 unilateral SCN from three VIP-Cre and four AVP-Cre mice). **f,** VIP Hub neurons had higher Reverse Pagerank compared to other cells, acting as generators of circadian synchronization signals (one-way ANOVA with Tukey’s post-hoc comparisons; *n* = 6 and 8 unilateral SCN from three VIP-Cre and four AVP-Cre mice). D, Dorsal; V, Ventral; M, Medial; L, Lateral; 3V, 3rd Ventricle; OC, Optic Chiasm.

We also found populations of SCN bridge and sink cells. Although both VL and DM cells received similar numbers of afferents (from 4.1 ± 0.7% vs. 4.2 ± 0.9% of the network, VL vs. DM, mean ± SD, *n* = 14 unilateral SCN), DM cells had higher Closeness_in-degree_ (0.3 ± 0.12 vs. 0.4 ± 0.07, VL vs. DM; Fig. 3a and Supplementary Fig. 13), i.e. they were topologically positioned to receive circadian signals faster. Projections from VL cells mostly terminated at the interface with the DM module where another population of cells sent and received projections from both modules. These bridge cells exhibited high Betweenness centrality and were well positioned to mediate circadian transmission across the dorsoventral SCN (Fig. 3a and Supplementary Fig. 13). In contrast, DM cells appear to serve as sink cells which received VL and DM inputs and projected primarily to other DM cells, resulting in high Pagerank values (0.02 ± 0.008 vs. 0.07 ± 0.02, VL vs. DM; Fig. 3a and Supplementary Fig. 13). Altogether, based on their functional connectomes, we identified four SCN cell types belonging either to the DM and VL modules, and posit that circadian signals critical to synchronize the network are generated and disseminated by two types of VL hub cells (with high Reverse PageRank and Out-degree) and propagated to the DM sink cells through bridge cells in the central SCN.

Based on their anatomical location, we hypothesized that hub cells in the VL module include SCN VIP neurons. To test this, we inferred connectivity from published SCN explant recordings^52^ similar to ours, but with VIP and AVP neurons identified (264 VIP and 572 AVP neurons among 1472 neurons recorded in three VIP-Cre and four Avp-Cre SCN explants; Fig. 3b). The mean degree (3.3 ± 1.3% vs. 3.6 ± 0.3%, Shan et al vs. this study, mean ± SD, *n* = 6 VIP-Cre and 8 AVP-Cre unilateral SCN, unpaired Student’s t-test, *p* = 0.6), mean path lengths (2.2 ± 0.4 vs. 3.3 ± 0.4, unpaired Student’s t-test, *p* = 0.69), and clustering coefficients (0.17 ± 0.07 vs. 0.19 ± 0.03, unpaired Student’s t-test*, p* = 0.7) of these SCN aligned with our data. We found that VIP neurons sent approximately three times more projections than AVP neurons, and some VIP neurons had notably more efferents (Fig. 3c). Unsupervised hierarchical clustering of VIP neurons by their out-degree consistently revealed two groups in each SCN (Fig. 3d and Supplementary Fig. 15a). Independently, k-means clustering similarly identified two VIP cell-groups with more than 90% agreement (mean Rand Index = 0.91). One subset of VIP neurons (31.7 ± 9.1%), termed the VIP hub cells, sent significantly more projections to 11.2 ± 0.9% of the network (had high Out-degree, 2,240 ± 175 projections/cell) and high Reverse Pagerank values compared to other non-hub VIP and AVP neurons (Fig. 3f). Interestingly, these VIP hubs also projected distally and sent about two times longer projections compared to other cells of the network (mean projection distance: 0.3 ± 0.06 vs. 0.18 ± 0.05, VIP vs. other cells; Supplementary Fig. 15b). We therefore conclude that a subset of VIP neurons are hubs that generate and broadcast circadian signals across the SCN through their long-range and highly influential connectivity.

### A Subset of Distinctly Connected VIP Hub Neurons Drive SCN Intercellular Synchrony

To test the roles of these different cell types in SCN function, we simulated the experimentally inferred SCN networks, ablated cells based on their centrality and measured their ability to synchronize or sustain circadian rhythms (Fig. 4a, left panel; see Methods; *n* = 12 unilateral SCN with 10 simulations each, one SCN topology could not be simulated to synchronize within our parameter space). Consistent with experimental results, intact SCN networks (Control) synchronized their PER2 rhythms, with increasing coherence over six simulated days (Fig. 4a and Supplementary Fig. 16a). Intriguingly, the synchronization rates and phase relationships of the dorsomedial and ventrolateral SCN cells highly correlated between in silico and experimentally recorded explants, indicating that cell-cell connectivity architecture strongly dictates circadian synchronization patterns in space and time (Fig. 4b).

**Fig. 4.**
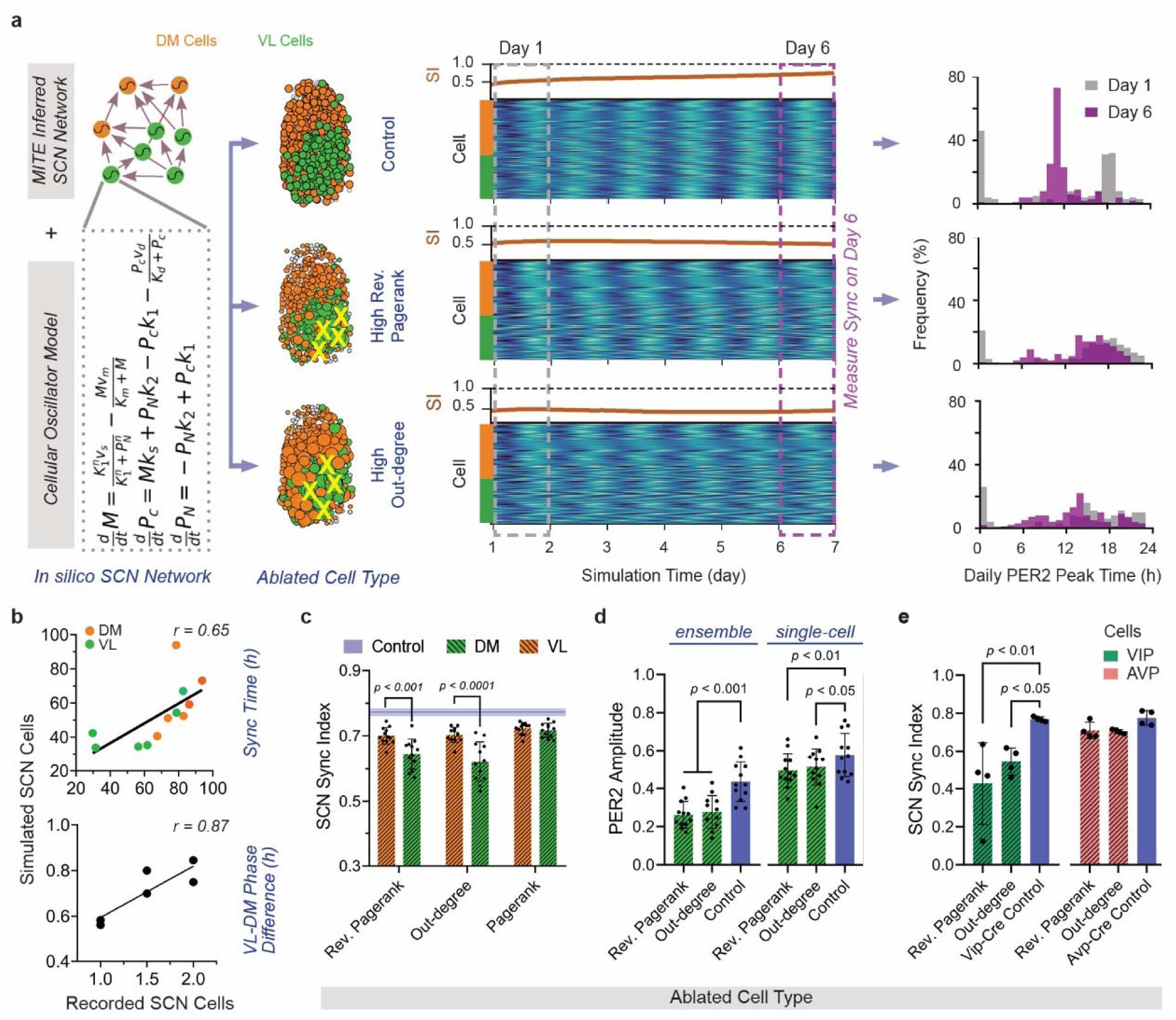
Circadian synchronization of the SCN requires VIP hub neurons. **a,** To test the roles of different cell types in synchronizing circadian cells of the SCN, we simulated the ability of in silico SCN networks with connectivity inferred experimentally from unilateral PER2::LUC recordings to recover from a desynchronized state (left). Raster plots from three representative simulations show how PER2 expression of DM (orange) and VL (green) cells changed over 6 days (middle). Ablating (yellow X) 30% VL cells with high Reverse Pagerank (signal generators) or Out-degree (broadcasters) impaired synchronization, as shown by their low sync index (SI, brown line) on day 6 compared to the Control SCN with no cells ablated. Similarly, PER2 peak-time distributions (right) on day 1 (grey dashed box and histograms) vs. day 6 (purple dashed box and histograms) showed spontaneous synchronization in the intact control SCN, but not when key cells were deleted. **b,** Time required by the DM and VL modules in simulated control SCN networks to reach synchrony (top) and their steady state circadian PER2 phase relationships (VL-DM phase difference, bottom) highly correlated with their behavior in recorded SCN explants (*r*, Pearson correlation coefficient, *n* = 6 SCN, reliable fit was not obtained for one SCN). Data for simulations was averaged across left and right SCN to facilitate comparisons **c,** SCN networks with 30% highly central (Reverse Pagerank or Out-degree) VL cells ablated (green) had reduced synchrony 6 days after ablation compared to DM cell ablations (orange). Blue line and shaded region are mean ± SD for control (no cells ablated, two-way ANOVA with Tukey’s post-hoc comparisons, *n =* 12 unilateral SCN) **d,** Ablating 30% of the highly central VL cells reduced daily PER2 amplitudes of individual cells and their ensemble rhythms measured 6 days after ablation. **e**, Ablating 30% of VIP cells (green striped bar) with high Reverse Pagerank or Out-degree reduced SCN synchronization compared to ablation of AVP cells (coral striped bar, two-way ANOVA with Tukey’s post-hoc comparisons, *n* = 4 unilateral). D, Dorsal; V, Ventral; M, Medial; L, Lateral; 3V, 3rd Ventricle; OC, Optic Chiasm.

Ablating cells based on their centrality dose-dependently reduced the ability of the in silico SCN to self-synchronize (synchronizability). Specifically, we tested the effects of deleting 10-30% of cells with highest centrality values and found that deleting the VL hub cells with high Reverse Pagerank or Out-degree (generators or broadcasters) had the greatest impact on synchronizability (Supplementary Fig. 16b,c). Ablating 30% of the cells with top Reverse Pagerank or Out-degree in the VL module reduced SCN synchrony by up to 20% (day 6 sync index: 0.78 ± 0.04 vs. 0.64 ± 0.05 vs. 0.62 ± 0.06, Control vs. Rev. Pagerank vs. Out-degree VL cells ablated, mean ± SD, *n* = 12 unilateral SCN with 10 simulations each; Fig. 4c). This globally reduced coherence among both VL and DM cells (Supplementary Fig. 16d,e), while halving the amplitude of daily PER2 rhythms (0.43 ± 0.09 vs. 0.26 ± 0.04; Fig. 4d). In contrast, ablating the top Reverse Pagerank or Out-degree cells in the DM module did not impact SCN synchrony (day 6 sync index: 0.78 ± 0.04 vs. 0.70 ± 0.02 vs. 0.70 ± 0.02, Control vs. Rev. Pagerank vs. Out-degree DM cells ablated; Fig. 4c). This further demonstrates that VL cells propagate information globally across the SCN while DM cells tend to signal locally (Supplementary Fig. 16e). We next similarly deleted cells in fully synchronized SCN (on day 6 rather than 0 of the simulation) and found that, in general, loss of 30% hubs (generators and broadcasters) in the VL module or bridge cells decreased network’s ability to sustain synchrony (Supplementary Fig. 17).

Using the same strategy on networks simulated with the connectivity maps derived from recordings of four unilateral VIP-Cre and AVP-Cre SCN each, we found that loss of high Reverse Pagerank or Out-degree VIP hubs reduced the sync index drastically by up to 44% (day 10 sync index: 0.77 ± 0.01 vs. 0.42 ± 0.21 vs. 0.54 ± 0.07, VIP-Cre Control vs. Rev. Pagerank vs. Out-degree VIP cells ablated, mean ± SD, *n* = 4 unilateral SCN/genotype with 10 simulations each; Fig. 4e). These simulations predict that while AVP neurons act primarily as outputs of the SCN network, VIP Hub cells, a third of VIP neurons, are critical to initiate and to sustain high amplitude, synchronous circadian rhythms in the SCN.

## Discussion

Mapping structure-function relationships in neural networks is fundamental to understanding how brains compute^17–20^. We report five, independent, in vitro and in silico tests indicating that MITE faithfully maps directed functional connections among SCN cells. Cellular connectivity maps inferred for 9011 SCN cells across 17 mice (including 264 VIP and 572 AVP neurons) revealed that connection patterns are organized by SCN region, reminiscent of the anatomically defined core and shell^8,21,24,108^. While the AVP and other cells in the dorsomedial SCN appear to act as sinks where circadian signaling converges, ventrolateral cells constituting a subset of hub VIP neurons (about 30% VIP cells) have critical efferent patterns that synchronize circadian cells throughout the SCN. This topology suffices to drive circadian synchronization and the daily wave of clock gene expression from dorsal to ventral SCN.

We found directed, sparse and efficiently coupled SCN cells connected as a small-world network, consistent with findings from inferred undirected connections^67^ and with the principle that brain networks minimize wiring costs while maximizing functional efficiency^109^. Specifically, MITE estimates an average SCN cell communicates directly with 719 ± 72 other cells yet, can signal to anywhere in the bilateral SCN through just 3-4 intermediate cells. This is remarkably consistent with recent estimates of 4 intermediate cells based on one anatomically reconstructed micro-network of 45 SCN neurons^63^. In agreement with predictions from theory^69^, histology^110^ and imaging^63,111^, we identified long-range cellular connections that underlie SCN synchronization, especially arising in the ventrolateral SCN. The cells with long range projections up to half-way across the SCN included the VIP hub neurons. Given the electron microscopy estimates that each SCN neuron makes about 450 synaptic contacts^24,63,112^, our results suggest non-synaptic volumetric transmission largely underlies efficient and long-distance circadian information transmission across the SCN while the short time-scale synaptic communication, like GABAergic signaling modulates the stability of SCN rhythms^68,113^.

Previously, undirected connections inferred from PER2 expression recordings revealed that all SCN cells had similar connectivity (degree, defined as number of connections/cell, were exponential distributed)^67^, whereas directed connections inferred from the same data suggested that SCN could contain hub cells with large number of efferents (scale-free degree distribution)^89^. We find that ventral SCN cells have more efferents and properties intermediate to scale-free and exponential connectivity (both in-coming and out-going connections per cell were best fit by Weibull curves^92^), consistent with anatomical evidence for heterogeneity in the number of dendrites per SCN cell^63^. We conclude that the heterogeneity of SCN connections allows for sparse and efficient wiring with spatially distinct long-range interactions that drive intercellular circadian synchrony^69,91^. Given that modelling predicts SCN circadian dynamics and output as attributable to cellular heterogeneity^114,115^, future studies to manipulate arborization in specific SCN cell-types or regions will help understand the role of heterogenous connectivity in SCN function. Alternatively, given that SCN undergoes sex-dependent spatiotemporal synaptogenesis and the dorsoventral phase-wave develops postnatally^116^, MITE connectivity maps inferred from SCN of neonatal and postnatal mice can reveal the sexually dimorphic functional roles of SCN connectivity topology.

Based on their relative density of connections, we identified two reproducible cellular modules: a small, densely connected ventrolateral module that sent over six times more projections, both locally and globally, compared to the larger and sparsely connected dorsomedial module, which mostly projected locally. Notably, this modular architecture was not detected in explants that failed to synchronize, indicating the two modules are strongly linked to SCN synchronizability. These findings, remarkably similar to the core and shell SCN regions previously defined by their anatomical, light response, and gene expression patterns^117–124^, further support the conclusions that MITE faithfully uncovers functional roles of anatomical connections and aids structure-function mapping in neural networks. Although over 40 SCN cell-types have been described by their anatomical, transcriptomic, histochemical, electrical, and light-response properties^8,21–31,63^, based on their functional interactions and simulations of experimentally inferred networks, we predict that four cell classes within the dorsal and ventral modules determine SCN function: Two types of Hub (circadian signal Generators and Broadcasters), Bridge, and Sink cells. While VIP is considered a critical synchronizer of SCN cells^29,39–47^, some recent findings have challenged this, placing AVP neurons as crucial for SCN synchronization^48–54^. We reveal that approximately one-third VIP neurons, based on their inferred connectivity and in silico ablation effects, function as SCN synchronization hubs. These VIP hub neurons, at the top of the signaling hierarchy, generate and disseminate synchronization signals through extensive and long-range projections. In contrast, AVP neurons appear dispensable, aligning with the loss of behavioral and body temperature rhythms observed in VIP-ablated but not AVP-ablated mice—an effect that seems age-dependent and warrants further investigation^29,125^. Based on recent findings that functioning clocks only in a subset of VIP neurons, the NMS-expressing VIP (VIP_NMS_), but not VIP_GRP_ neurons, are critical for maintaining circadian body temperature and behavioral rhythms^29^ and that these neurons exhibit photoperiod-induced neurotransmitter switching^126^, we hypothesize the VIP hubs critical for SCN synchrony are the VIP_NMS_ neurons. Notably, Bridge cells positioned between the dorsal and ventral SCN modules, resemble the Prokineticin 2 (PROK2) and possibly, GRP neurons. Since GRP expression is unaffected by photoperiod^126,127^, we speculate that, by their network positioning as bridges, and their daily and seasonal changes in signaling^25,127^, PROK2 cells modulate information flow to couple the dorsal and ventral SCN and reorganize the two regions across seasons. Studies to test the impact of ablating these cell populations on SCN coherence and entrainment to different day-lengths will clarify the roles of these functional cell-types.

Interestingly, circadian dynamics of cells when connected in silico using MITE inferred SCN maps mirrored their experimentally recorded explants. Ventrolateral cells not only synchronized faster than dorsomedial SCN in silico, their PER2 expression recapitulated the dorsoventral phase-wave. These exciting findings, consistent with evidence that cell-intrinsic differences do not drive the daily PER2 wave^16,119,128^, demonstrate that emergent properties like spatial and temporal synchronization among SCN cells are sculpted by their connectivity architecture. Finally, while the functional connectome complements anatomical and transcriptomic classification of SCN cell populations, it enhances our understanding of the roles of these cell types in network behavior and circadian time computation.

## Methods

### In Silico Network Simulation

To compare accuracy of different network inference methods, we generated networks of 600 nodes to approximate the number of cells recorded per SCN in our experiments. Each node was modeled as a cellular circadian oscillator using ordinary differential equations, describing the transcription of the PERIOD2 gene mRNA (*M_i_*) and production of cytosolic (*P_C_*) and nuclear ^82,129^ PER2 protein (*P_N_*) in a single cell. Cellular heterogeneity was introduced by varying the PER2 mRNA degradation rate (*v_m,i_*) between 0.345 -0.395, sufficient to generate cellular periods of 23.4 ± 0.7 h (mean ± SD), as observed experimentally^11^. All other model parameters were similar to prior studies (Supplementary Table 2)^82^. The intercellular coupling influences the mRNA transcription rate (*v_s,i_*) of cell *i*, which is defined as 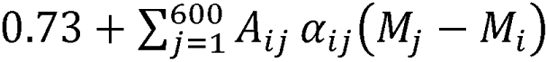, where *α_ij_* denotes the strength of the input from cell *j* to cell *i* and *A_ij_* = 0 or 1, indicating absence or presence of connection between the cells *i* and *j*. Coupling strengths were selected such that the network synchronizes within 4-5 days to mimic experimental observations.

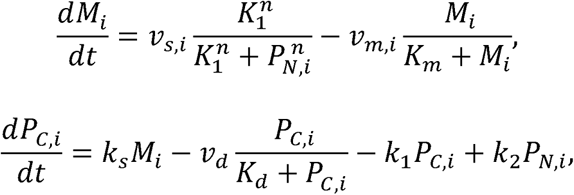

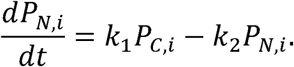

We modeled small-world, scale-free, and mixed (a combination of small-world and scale-free) networks using Python NetworkX package^130^. The directed scale-free networks were generated using the ‘scale_free_graph’ function, with parameters α and β varying in the ranges [0.6, 0.69] and [0.26, 0.35], respectively, and y set to 0.05, ensuring that *α* + *β* + *γ* = 1. Here, *α* represents the probability of adding a new node connected to an existing node selected randomly based on its in-degree distribution (distribution of incoming connections), *β* prepresents the probability of adding an edge between two existing nodes (with one node chosen randomly based on its in-degree and the other based on its out-degree–number of outgoing connections), and *γ* represents the probability of adding a new node connected to an existing node selected randomly based on its out-degree distribution. For small-world networks, undirected networks were first generated using the ‘connected_watts_strogatz_graph’ function, with the probability of rewiring each edge varying uniformly between [0.1, 0.25] and the mean degree varying between [4, 7]. We randomly deleted 100 edges from the connectivity matrix to convert the network into a directed one. Mixed networks were generated by combining outgoing connections (connectivity matrices) from both small-world and scale-free topologies. Each network was simulated 10 times varying the initial conditions and coupling strengths.

To approximate the spontaneous synchronization dynamics of desynchronized SCN neurons after tetrodotoxin (TTX) removal, we simulated in silico networks from random initial conditions and recorded their PER2 mRNA at 1-hour intervals for 6 days, as in our experimental SCN recordings. We quantified coherence among circadian cells as the Synchrony Index (sync index, 0=circadian cells peak at random times relative to each other; 1=cells all peak at the same time) using the first-order Kuramoto order parameter by vectorial averaging of instantaneous phases on the complex plane^131^.

### Comparing MITE to Other Network Inference Methods on In Silico Data

By pairwise comparisons of the simulated PER2 traces of all nodes, we generated an interaction score matrix for all possible cell-cell interactions in the network. We reasoned that if two cells couple, their PER2 dynamics should co-evolve as the network synchronizes, resulting in high interaction scores, whereas disconnected cells should exhibit lower scores. We inferred interaction scores using two categories of methods, each with three different statistics. The first category, undirected methods, included Dynamic Time Warping (DTW)^132^, Maximal Information Coefficient (MIC)^77^ and Mutual Information (MI)^78^, which identified cell-cell interactions without considering interaction direction. The second category, directed methods, inferred interaction direction and included the Algorithm for Revealing Network Interactions (ARNI)^79^, Granger Causality (GC)^80^ and Transfer Entropy (TE)^81^.

DTW analysis was implemented using the ‘distance_matrix_fast’ function in DTAIDistance Python package^133^. Since DTW measures dissimilarity between time series (higher scores indicate weakly connected cells), the DTW scores were inverted for further analysis. MI was estimated using ‘mutual_info’ function in PyInform^134^ python library and MIC analysis using MICtools for Docker^135^, with parameters previously used for analyzing similar datasets of spontaneously synchronizing SCN cells^67^. Unlike Abel et al.^67^, who used heuristic APPROX-MIC scores, MICtools computes MIC using a combination of MIC-based measures, MICe (a consistent estimator of the MIC population value) and TICe (total information coefficient)^135,136^, which have better bias/variance properties and are more computationally efficient.

ARNI was implemented in Matlab (The MathWorks Inc. (2022). MATLAB version: 9.13.0 (R2022b), Natick, Massachusetts: The MathWorks Inc. https://www.mathworks.com) using the author-provided code^79^. GC and TE analyses were performed using ‘grangercausalitytests’ and ‘transfer_entropy’ functions in python statsmodels^137^ and PyInform^134^ libraries respectively. The parameters maxlag (for GC) and history length (for TE) that decides the number of previous time points considered to compare two time series, was set to 1 because our in silico networks did not model delayed coupling. We computed univariate TE (considers two cells at a time), which is equally accurate (for short time-series) as the computationally intensive multivariate TE^74^. We inverted the resultant TE values to represent directed interactions among synchronizing oscillators. To generate MITE connectivity matrix (Mutual Information & Transfer Entropy), we scaled the MI and TE scores using min-max scaling to range between 0 and 1, multiplied the two, and z-transformed them to produce MITE scores for each cell-cell interaction.

We computed the Area Under the Receiver Operating Characteristic (AUROC) of MITE connectivity matrix by comparing to the true connectivity matrix using the scikit-learn python library^138^. For each network, we computed the AUROC across 10 simulations and averaged for statistical analysis. All analyses were performed on a Dell Precision 5820 Tower X-Series (Intel(R) Core(TM) i9-10920X CPU @ 3.50GHz, 64.0 GB RAM, NVIDIA GeForce RTX 3090) or a Dell OptiPlex 7000 (12th Gen Intel(R) Core(TM) i7-12700 @ 2.10 GHz, 32 GB RAM, Intel(R) UHD Graphics 770).

### Animals and Housing Conditions

Homozygous *PERIOD2::Luciferase* (PER2::LUC) knock-in mice (founders generously provided by J.S. Takahashi, UTSW) were housed under a 12-hour light:12-hour dark cycle maintained at constant temperature of 25^0^C with food and water ad libitum in the Danforth Animal Facility, Washington University in St. Louis. All animal procedures were approved by the Animal Care and Use Committee at the university and adhered to NIH guidelines.

### Single Cell Bioluminescence Recording of PER2::LUC SCN Explants

Coronal brain slices (300 μm thick) were obtained from neonatal (P4-P7) PER2::LUC mice using a vibratome (EMS). The slices were immediately placed in chilled Hanks’ balanced salt solution (HBSS) supplemented with 0.01 M HEPES, 100 U/ml Penicillin, and 4 mM NaHCO3. The SCN region was carefully dissected from these slices using scalpels and cultured on 0.4 mm membrane inserts (Millicell-CM, Millipore) with 1 ml of HEPES-buffered air-DMEM (1.2 ml), supplemented with 10% newborn calf serum (Invitrogen) and 0.1 mM beetle luciferin (Biosynth). Each SCN sample was recorded using an ultrasensitive CCD camera (Andor Ixon or Ikon; 1 x 1 binning, 1-hour exposures) at 36°C. The recordings were conducted on an inverted microscope (Nikon TE2000 or Leica DMi8) equipped with a 20X objective and a 0.5X coupler, within a glass-bottom, sealed Petri dish containing media supplemented with 0.1 mM beetle luciferin (Biosynth) inside an In Vivo Scientific incubator, as described previously^67,102,139^.

Bioluminescence recording was started one day after the slice preparation and following 4-5 days of baseline recording, SCN explants were treated with 1.5 µM or 3µM tetrodotoxin (TTX, Sigma) to block spiking and cell-cell communication among SCN neurons for 6 days. Three full-volume exchanges of fresh medium were done and the washed SCN was recorded for at least another 6 days to monitor the resynchronization of cellular rhythms following the re-establishment of cell-cell communication. To identify and track cellular PER2 bioluminescence, we employed a custom Python code that detected cells in each frame using a Difference of Gaussian blob detector^67^. The pixel intensities of the identified cells were measured over the course of the recording to extract PER2 time series traces for further analysis. Data from first 12 hours of recording were not considered due to artifacts arising from TTX-wash or culture medium change.

### Circadian Rhythm Analysis of PER2 Gene Expression

We measured periodicity of PER2 traces of all identified SCN cells with Lomb-Scargle implemented in MetaCycle^140^ and identified significantly circadian cells (BH.Q < 0.01 within a period range of 18 to 30 hours). We detrended and smoothed PER2 traces from circadian cells to compute instantaneous phase with continuous wavelet transformation and the Synchrony Index among cells as the first-order Kuramoto order parameter^131^ implemented in pyBOAT^141^.

To analyze time taken by dorsal and ventral modules to synchronize, PER2 signals were averaged across cells within the module, and daily PER2 peak-to-peak periods were estimated using peak finding and fit to a Gompertz curve^142^ in SciPy^143^ python library. The curve fit provided the time rate of period change and we defined time to synchronize as the time after which the rate of period change was less than 0.1 h for the rest of the recording (0.01 h was used for in silico cells).

### Inferring SCN Cell-Cell Connectivity Using MITE

Detrended PER2 bioluminescence traces recorded from synchronizing SCN cells after TTX removal were used to compute z-transformed pairwise MITE interaction scores (MITE connectivity matrix) as described for simulated networks, with the exception that history length was set to 4 hours to incorporate delayed cellular coupling (simulated networks were analyzed with history length of 1 as we did not model delayed interactions). The MITE connectivity matrix was then thresholded using an empirically determined cutoff of 2.07 (1.03 was used for Vip-Cre and Avp-Cre SCN). This resulted in a MITE binary connectivity matrix with scores below this threshold considered as false cell-cell connections and those above as true connections. The binary connectivity matrix was used for further network analysis. Self-loops (connections from and to the same cell) were not considered for analysis. PER2 traces from AVP-Cre and VIP-Cre SCN reported in Shan et al^52^ were analyzed similar to our recordings. To ensure accurate estimation, cells with < 160 h of data or with > 1 h gaps in data were not considered for analysis.

To determine the cutoff MITE score, we pooled PER2 traces from cells across four randomly chosen SCN constituting the Training set and estimated interaction scores of the impossible false connections identified between SCN of different mice (3081 cells with 9.49 million pairwise comparisons). We empirically determined the threshold z-score (2.07) that minimizes the probability of detecting these ‘impossible between-SCN’ connections (false discovery rate < 0.05) while maximally retaining the possible within-SCN connections (hit-rate > 95%). We similarly tested whether the chosen threshold reliably identified connections in an independent Test set with 3 SCN (1994 cells with 3.97 million comparisons). Connections between left and right SCN of the same mouse were not considered as they cannot be categorized as impossible connections. Threshold score for AVP and VIP connectivity were estimated similarly by pooling cells from seven Avp-Cre and VIP-Cre SCN (1472 cells with 3.16 million comparisons).

### In Silico Cell Ablation

We compared circadian synchronization properties of in silico networks with targeted cell ablations. We simulated unilateral SCN networks generated using MITE binary connectivity matrices inferred from experimental recordings and simulated each SCN 10 times, varying the initial conditions and coupling strengths. All simulated SCN started in a desynchronized state on Day 1 and synchronized their PER2 rhythms by Day 6. We ablated cells with the top 10%, 20%, or 30% of specified centrality scores by deleting their incoming and outgoing connections and then repeated the simulations with the same initial conditions and coupling strengths as their corresponding unablated control SCN network. We measured sync index (SI) on Days 1 and 6 using the Kuramoto order parameter averaged over 24 hours. Separately, to investigate the effect of cell ablation on sustaining synchrony, we repeated the simulations, deleted cells on Day 6 and similarly measured SI on Day 10 averaged over 24 hours.

### SCN Network and Spatial Connectivity Analysis

We used the inferred MITE binary connectivity matrices to analyze SCN networks using Cytoscape^144^, Gephi^145^ and NetworkX packages^130^. We used author provided code to measure graphlet based network similarity (DGCD-13 and DGCD-129)^83^ as distances between each unilateral/bilateral SCN network and ten corresponding random networks with similar numbers of cells and connections, but with randomized connection patterns. We report the average DGCD distances from each SCN to 6 other SCN and with 10 corresponding random networks. We measured the small-worldness coefficient defined as the ratio of clustering coefficient (C) and path length (L) of the SCN and corresponding random networks (denoted by subscript r) with similar number of nodes and connections -[C/C_r_]/[L/L_r_][C/C_r_]/[L/L_r_][C/Cr]/[L/L_r_]^146^. Scale-free exponent (alpha) was estimated as the slope of power-law function (P_k_ ∼ k^-α^) fitted to the probability distribution (P_k_) of log-transformed out-degree (*k*, number of efferents).

To identify cellular communities within each unilateral SCN, we used NetworkX^130^ functions ‘greedy_modularity_communities’, ‘label_propagation_communities’, and ‘girvan_newman’. The identified communities were clustered using k-means and hierarchical clustering based on their mean out-degree using SciPy^143^ with latter parameters: linkage method=’ward’, distance metric=’euclidean’. Other cell-cell connectivity statistics were analyzed using custom python scripts. To assess spatial distribution of cells within each module, we normalized the cellular coordinate space (X,Y coordinates were min-max scaled using the cells in extrema) and used a combination of Kernel Density Estimation (‘gaussian_kde’ function in Scipy^143^) and local maxima detection (‘peak_local_max’ function in Scikit-image^147^) algorithms to identify the regions with high cell density. Histograms of cell density along the dorsal-ventral and medial-lateral axes were analyzed by binning cells in 10% and 20% bins respectively along the two axes. Similarly, to analyze projection distances, we used the normalized coordinate space and calculated the Euclidean distance of each cell to all its connecting cells. We used NetworkX functions ‘in_degree_centrality’, ‘out_degree_centrality’, ‘betweenness_centrality’, ‘closeness_centrality’ (measures closeness_in-degree_), and ‘pagerank’ to compute centrality scores for each cell. The closeness_out-degree_ and reverse pagerank centralities were analyzed using the same functions with transposed binary connectivity matrices to reverse the connection direction. Centrality values were min-max scaled to facilitate inter-SCN comparisons.

All statistical analysis were implemented in GraphPad Prism version 10.0.0 for Windows, GraphPad Software, Boston, Massachusetts USA, www.graphpad.com, and data were plotted using Prism, Matplotlib^148^ and Seaborn^149^.

## Data availability

Data presented in the present study are available upon request to the corresponding authors.

## Code availability

Code used for the present study are available upon request to the corresponding authors.

### Acknowledgements

The authors thank the members of the Herzog, Kiss and Li labs for valuable discussions and comments on the manuscript. This work was supported by National Institutes of Health Grants NS121161, NS139415, and GM157609.

## Author contributions

K.L.N, E.D.H, I.Z.K and J.-S.L conceptualized experiments and procured funding. K.L.N, B.S. and D.G designed experiments and collected data. K.L.N and B.S analyzed and interpreted data. K.L.N prepared the manuscript with inputs from other authors.

