## Supplementary Information for "The Functional Connectome Mediating Circadian Synchrony in the Suprachiasmatic Nucleus"

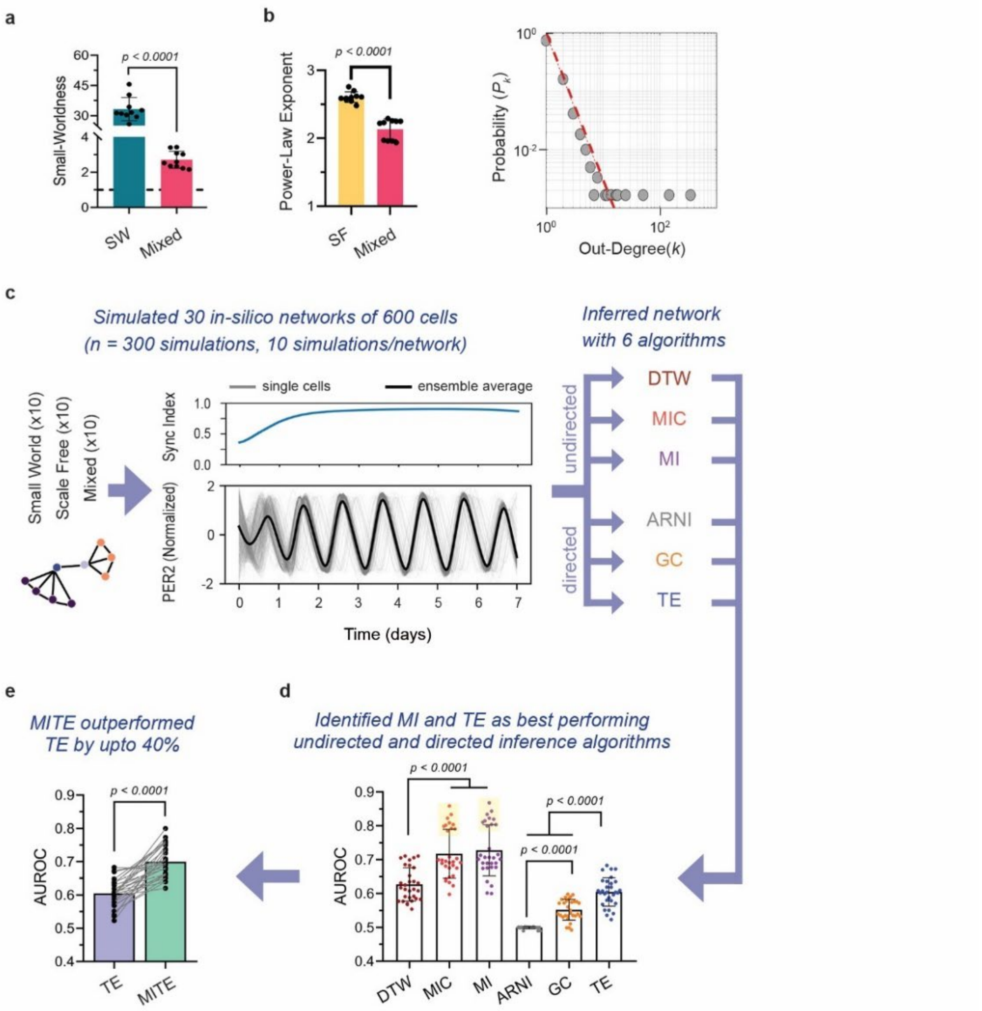


**Supplementary Fig. 1 | MITE outperforms other methods for inferring directed networks. a** and **b**, We tested MITE and other methods on in silico networks with SW topology (small-world) that displayed high small-worldness coefficient (threshold dashed line=1, **a**), SF networks (scale-free) that displayed high scale-free exponent–alpha, estimated from power-law fits to the log-transformed probability distribution $P_k$ of out-degree, k (number of efferents, gray circles), shown for a representative SF network and Mixed networks (mix of SW and SF) that displayed intermediate values of small-worldness and alpha (unpaired Student's t-test, $n$ = 10 networks per topology). **c,** Each of the 600 cells in each SCN model included a Period2 transcription-based circadian oscillator. Each simulation started from a unique desynchronized initial state. Shown for a representative simulation (middle), average PER2 (thick line) and synchrony index (measure of network coherence, blue line) calculated from the 600 cells (gray lines) that spontaneously synchronized similar to experimental recordings. We inferred the underlying connections of each simulated network using three undirected (right, DTW = Dynamic Time Warping, MIC = Maximal Information Coefficient, MI = Mutual Information) and directed (ARNI = Algorithm for Revealing Network Interactions, GC = Granger Causality, TE = Transfer Entropy) methods. **d,** Networks inferred by MI and TE had higher accuracy (Area Under the Receiver Operating Characteristic, AUROC, two-way ANOVA with Tukey's post-hoc comparisons, $n$ = 30 networks, dots show the average of ten simulations/network). MI and MIC were topology sensitive and inferred SW networks better than other topologies (yellow shaded box). **e,** Our algorithm, MITE (Mutual Information & Transfer Entropy) outperformed TE by up to 40% (paired Student's t-test, $n$ = 30 networks, dots show the average of ten simulations/network).

**Supplementary Figure 2** **Nikhil et al**

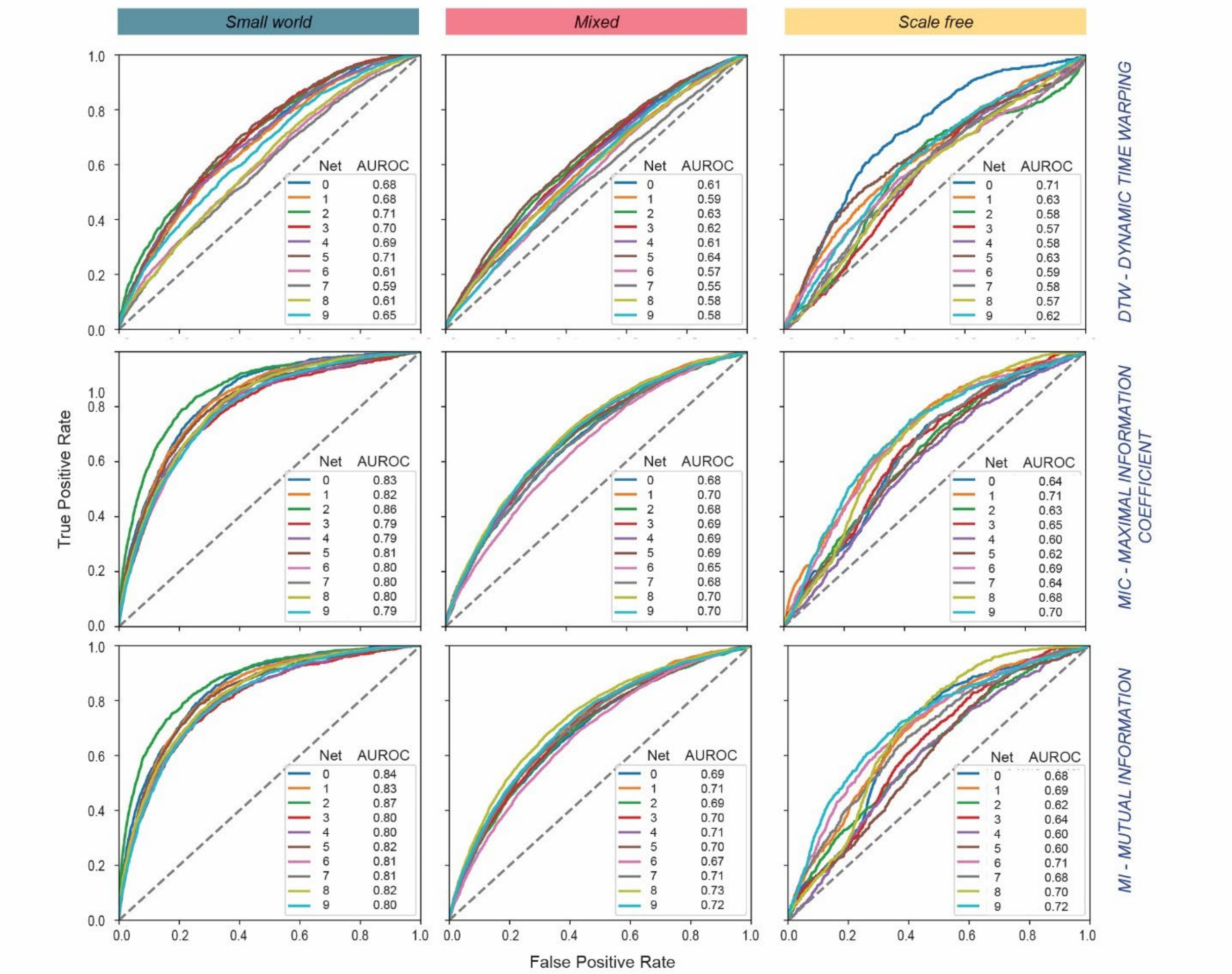


**Supplementary Fig. 2 | Performance of undirected cell-cell connectivity inference methods across 300 simulations of 30 in silico networks with different topologies.** Receiver Operating Characteristic curves (ROC) for 10 networks/topology (Net 0-9, colored lines) showing true and false positive rates of identified connections, with corresponding AUROC values (Area Under the ROC) for networks inferred by Dynamic Time Warping, Maximal Information Coefficient, and Mutual Information algorithms. Each ROC curve is the average of 10 simulations. Dashed gray line indicates AUC = 0.5, expected performance of a method that randomly guesses connections.

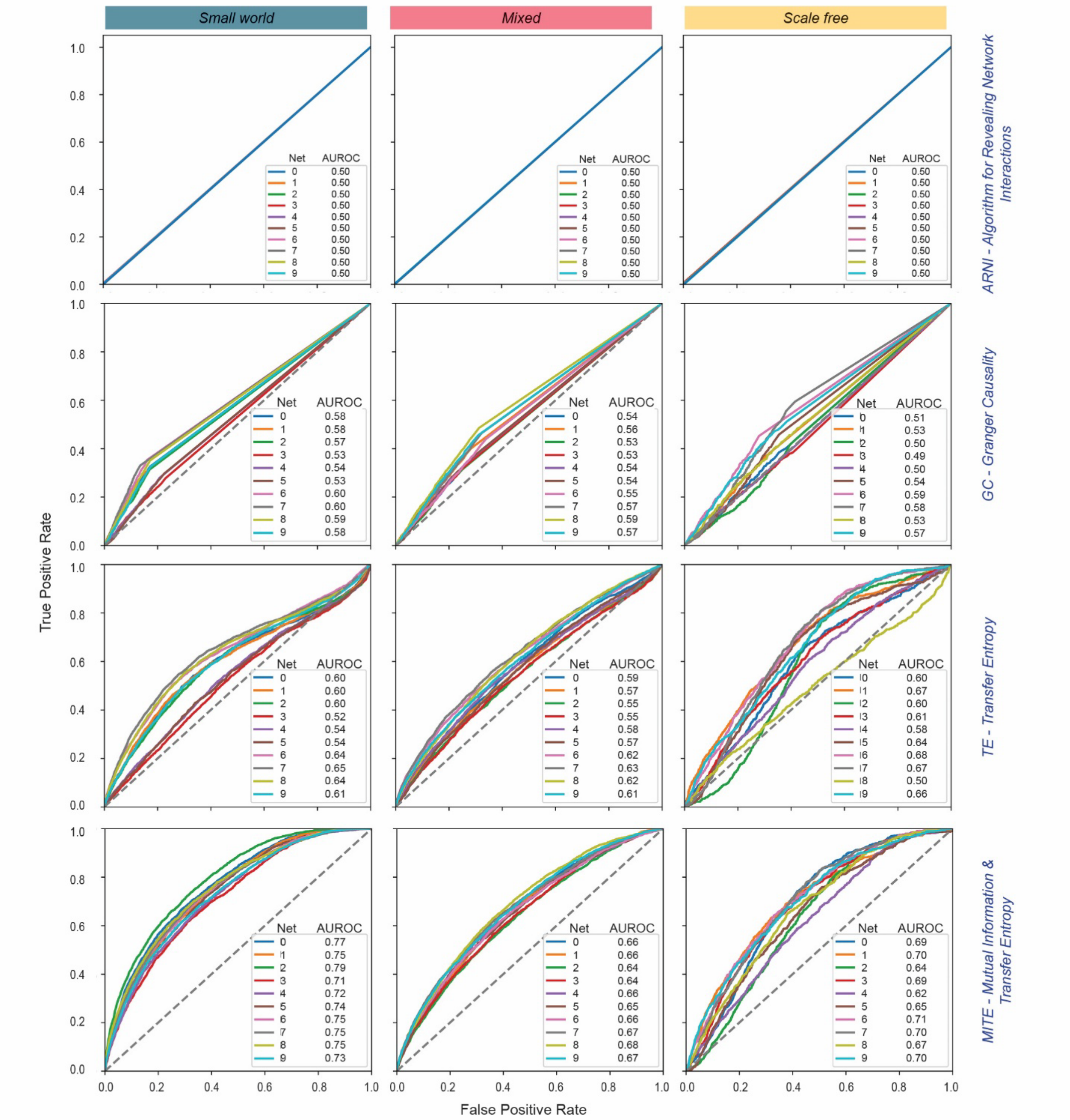


**Supplementary Fig. 3 | Performance of directed cell-cell connectivity inference methods across 300 simulations of 30 in silico networks with different topologies.** Receiver Operating Characteristic curves (ROC) for 10 networks/topology (Net 0-9, colored lines) showing true and false positive rates of identified connections, with corresponding AUROC values (Area Under the ROC) for networks inferred by Algorithm for Revealing Network Interactions, Granger Causality, Transfer Entropy and Mutual Information and Transfer Entropy.

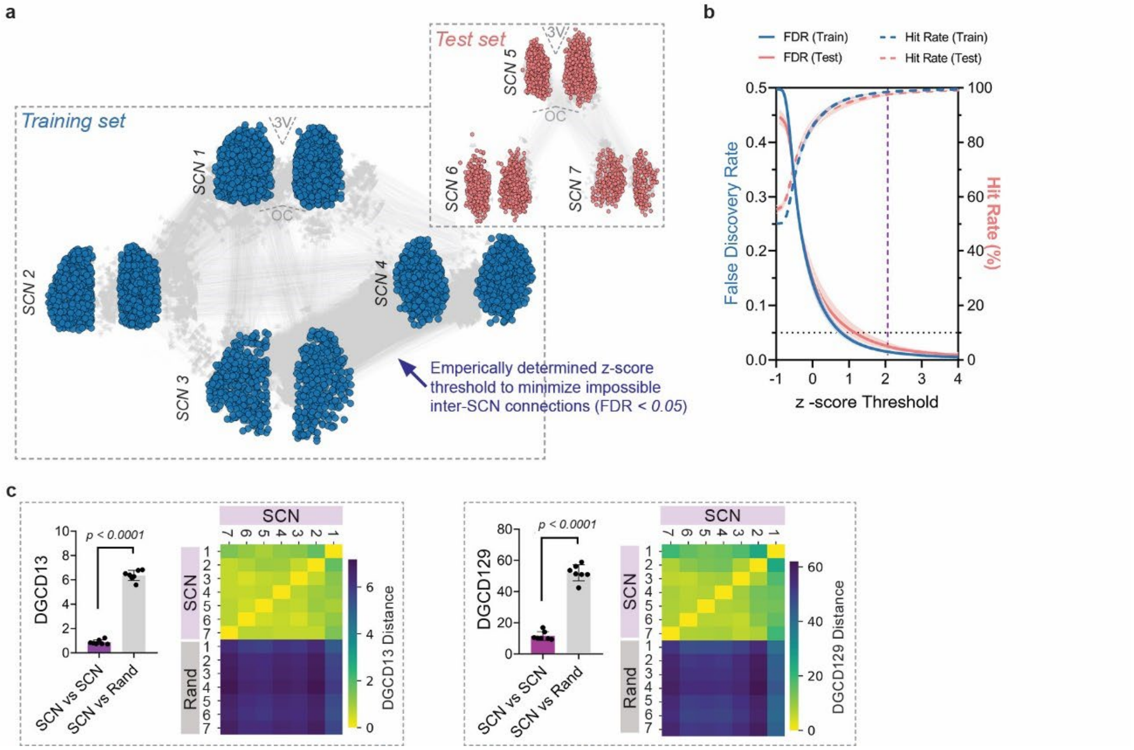


**Supplementary Fig. 4 | MITE infers SCN cell-cell interaction networks with high fidelity. a,** MITE analysis of PER2 expression data pooled from 3081 cells in four independent SCN (SCN1-4, Training Set, blue) included impossible false inter-SCN connections (grey lines). **b,** From 9.5 million pairwise comparisons across 3081 cells, we empirically determined the threshold z-score for MITE interaction scores (2.06, purple dashed line) to minimize impossible false inter-SCN connections and revealed networks with < 0.05 false discovery rates (probability of falsely identifying an impossible SCN connection, FDR, blue solid line) and > 95% hit rates (percentage of likely true intra-SCN connections among all identified connections, blue dashed line). The empirically chosen threshold robustly identified connections with < 0.05 FDR and > 95% hit rates (coral solid and dashed lines, respectively) when tested on PER2 traces of 1994 cells (3.97 million pairwise comparisons) in an independent Test set of three SCN (**a**-inset). **c,** Low Directed Graphlet Correlation Distance-13 (DGCD13, left) and DGCD-129 (right) values revealed that networks inferred by MITE were highly conserved across all seven SCN (SCN1-7) compared to random networks with similar numbers of nodes and connections (Rand1-7; unpaired Student's t-test). For SCN vs. SCN comparisons, values in bar plots are average distance of each SCN to six other SCN, and for SCN vs. Rand comparisons, values are average distance of each SCN to ten random networks. Heatmaps show the same data as pairwise SCN vs. SCN and SCN vs. Rand network comparisons. Error bars are SD. D, Dorsal; V, Ventral; M, Medial; L, Lateral; 3V, 3rd Ventricle; OC, Optic Chiasm.

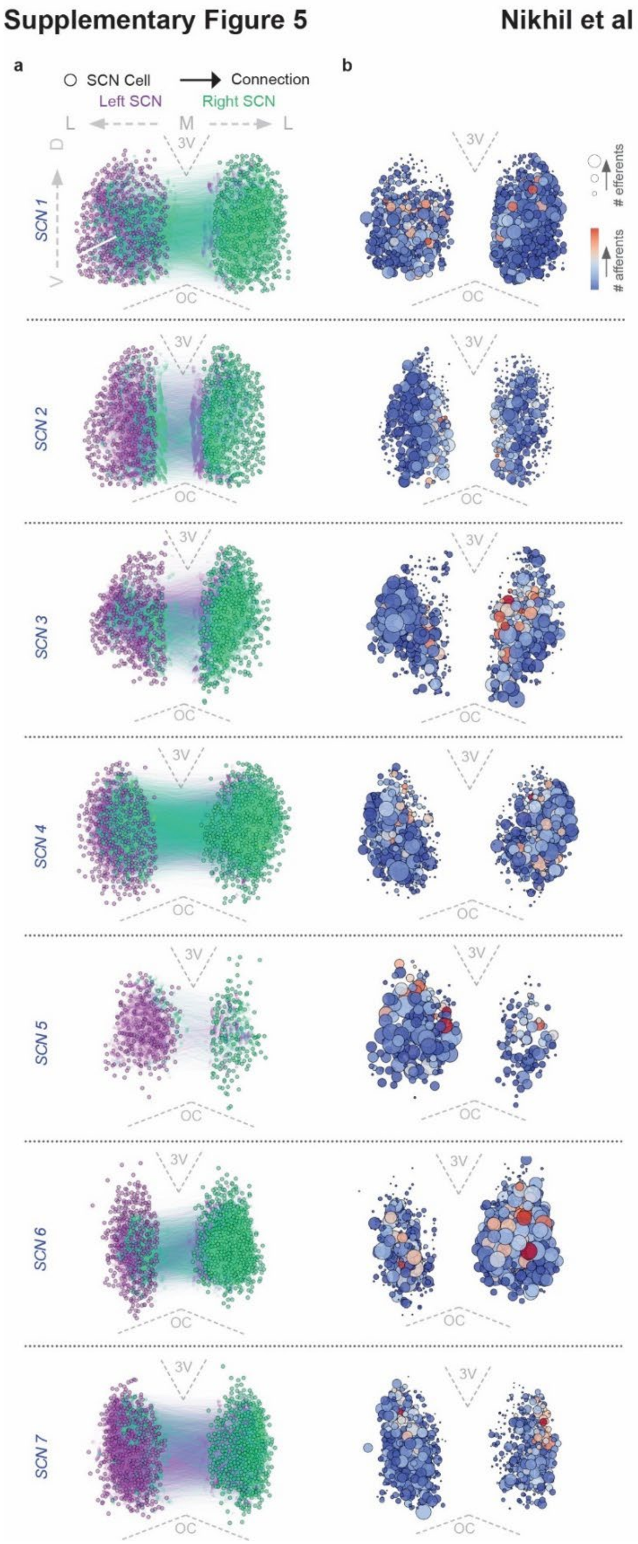


**Supplementary Fig. 5 | Bilateral and spatial connectivity patterns within the SCN. a,** Inferred maps of cell-cell connections (circles and lines) in seven SCN (SCN 1-7) show more ipsilateral connections (within left or right SCN) than contralateral projections. **b**, Ventrolateral SCN cells had more efferents (larger circles) while most SCN cells received similar numbers of afferents (blue saturation). D, Dorsal; V, Ventral; M, Medial; L, Lateral; 3V, third ventricle; OC, Optic Chiasm.

### Supplementary Figure 6



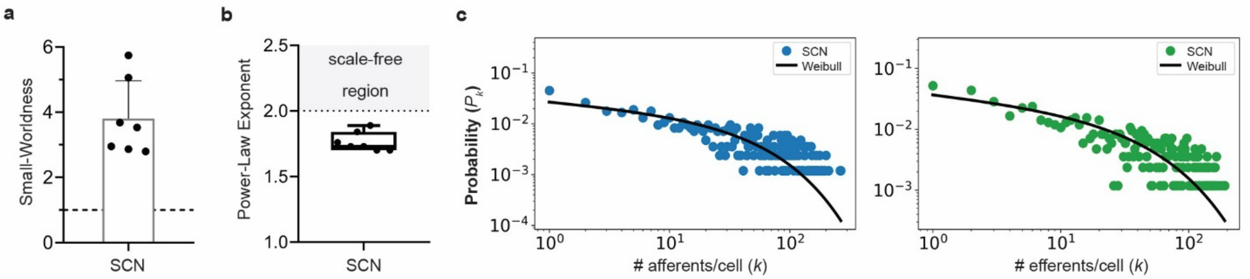


**Supplementary Fig. 6 | SCN connectivity has small-world topology with heterogeneously connected cells. a** and **b,** SCN cells were connected as small-world networks (small-worldness coefficient > 1, dashed line) and were not scale-free networks (power-law exponent < 2, estimated from power law fits to the probability distribution of out-going connections (number of efferents/cell). Error bars are SD. **c,** Probability distribution of incoming (number of afferents/cell, left) and out-going SCN connections (number of efferents/cell, right) were best fit by Weibull curves (solid lines) as estimated by Maximum Likelihood comparisons of multiple distributions, shown for a representative SCN (data plotted on log-log scale). Weibull distribution reveals that SCN cells have heterogenous connections with some exhibiting high connectivity.

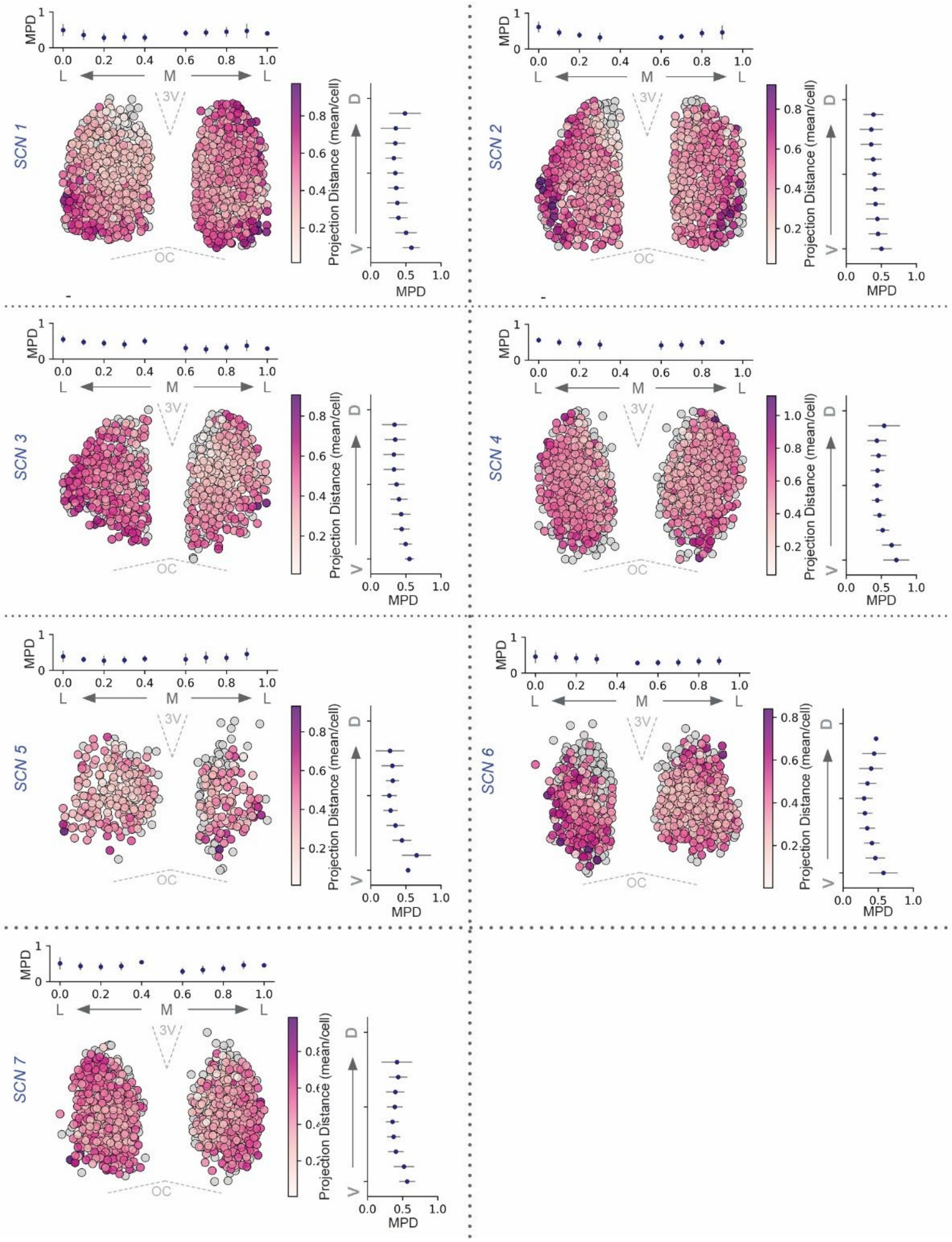


**Supplementary Fig. 7 | Ventrolateral SCN cells have longer efferents compared to dorsal cells. a,** Spatial maps of cellular projection distances in seven bilateral SCN. VL SCN cells project farthest based on cellular projection distances (mean Euclidian distance of a cell to all its targets, purple saturation), and mean projection distances (MPD, average projection distance of cells grouped in 10% bins) along the ventral-dorsal (right) and medial-lateral axes (top). Error bars are SD.

**Supplementary Figure 8** **Nikhil et al**

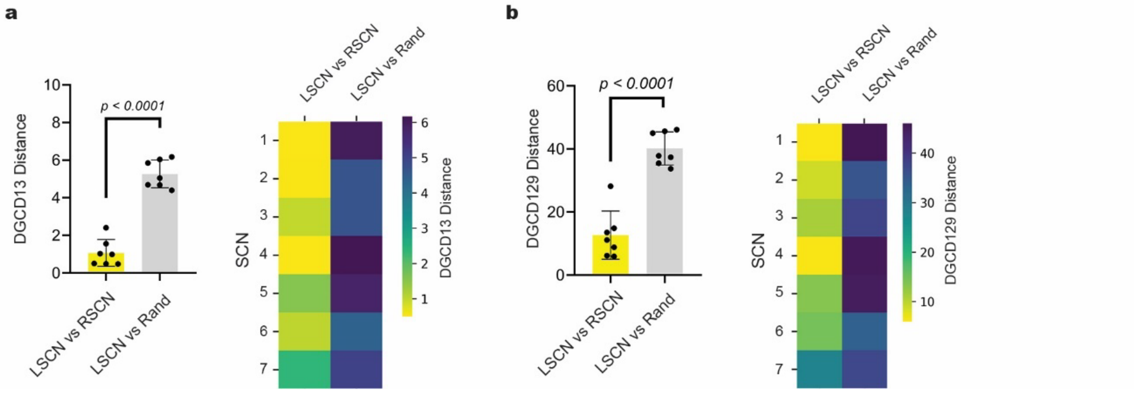


**Supplementary Fig. 8 | SCN networks are bilaterally similar.** Low Directed Graphlet Correlation Distance-13 (DGCD13, **a**) and DGCD-129 (**b**) values indicate that unilateral left and right SCN networks (LSCN and RSCN) are strikingly similar to each other but differ from random networks of similar size and connections (unpaired Student's t-test). For LSCN vs. RSCN comparisons, values in bar plots are average distances of each SCN to six other SCN, and for LSCN vs. Rand comparisons, values are average distances of each SCN to ten random networks. Heatmaps show the same data as pairwise LSCN vs. RSCN and LSCN vs. Random network comparisons. Error bars are SD. D, Dorsal; V, Ventral; M, Medial; L, Lateral; 3V, 3rd Ventricle; OC, Optic Chiasm.

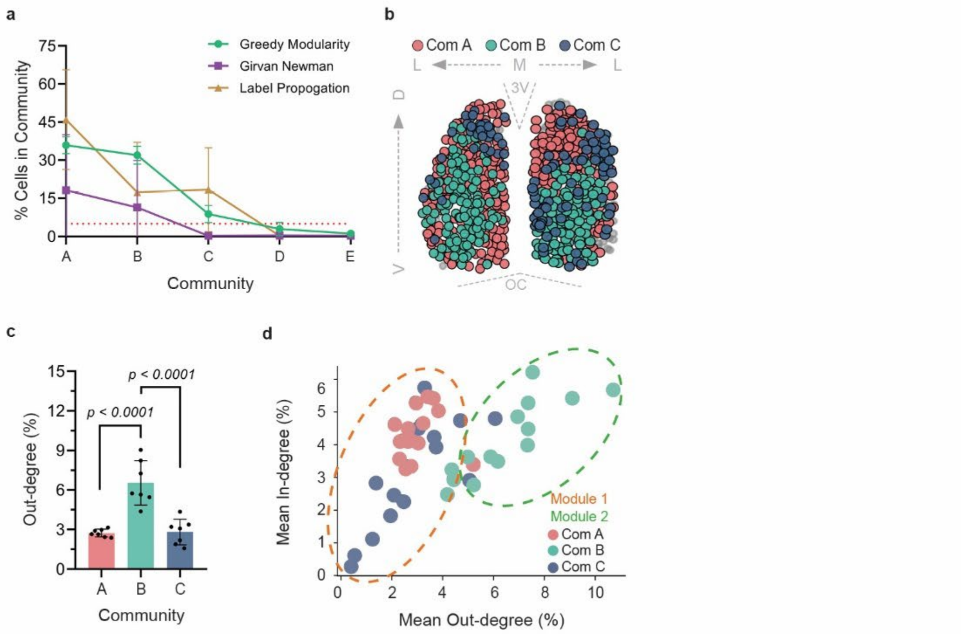


**Supplementary Fig. 9 | Cellular communities within the SCN clustered into two modules based on their connectivity**. **a,** Greedy Modularity, Girvan-Newman, and Label Propagation consistently revealed three communities (Com A-C) that, together comprised > 90% SCN cells (n=14 unilateral SCN). Any remaining communities had < 5% (red dotted line) cells. **b,** Spatial map of a representative bilateral SCN showing recorded cells (circles) in Comm A and C had similar spatial distributions. **c**) Communities A and C also had similar mean Out-degrees (average percent of the network to which a cell projects, A: 2.7 ± 0.3% , B: 6.5 ± 1.6%, C: 2.8 ± 0.9%, mean ± SD, $n$ = 7 SCN). **d**, k-means clustering by Out-degree of all 42 communities (filled circles) in 14 unilateral SCN grouped communities A and C together as a module in most SCN (module 1, coral dashed line) while community B was mostly an independent module 2 (cyan dashed line), shown using a scatter plot of their average Out-degree vs. In-degree (percent of the network from which a cell receives afferents). Error bars are SD. D, Dorsal; V, Ventral; M, Medial; L, Lateral; 3V, third ventricle; OC, Optic Chiasm.

#### Supplementary Figure 10 Nikhil et al

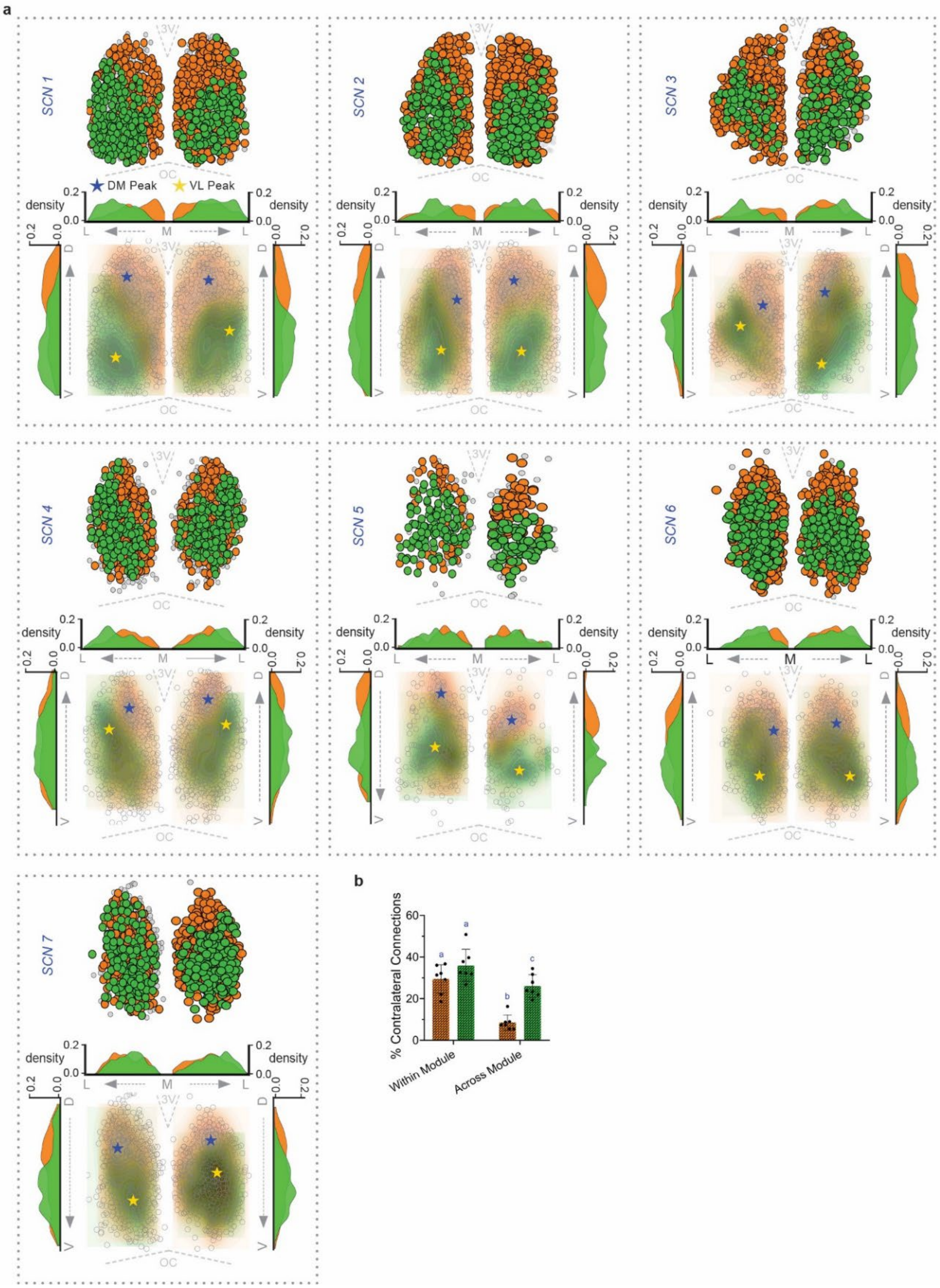


**Supplementary Fig. 10 | The SCN exhibits modular organization comprising asymmetrically connected dorsomedial and ventrolateral modules. a,** Top panel: Based on their connections, SCN cells clustered as Dorsomedial (DM, orange) and Ventrolateral modules (VL, green), with only a few excluded from either (grey circles), shown in a representative bilateral SCN, shown in all seven SCN (SCN 1-7).

Density contours (bottom panel) show spatial distribution of the DM and VL cells with their peak cell-density regions marked (blue and yellow stars). Histograms are cell-density along the dorsal-ventral (side histograms) and medial-lateral SCN (top histograms) axes. **b,** Contralateral projection patterns were similar to the ipsilateral connections. Of all contralateral connections, there were more projections across to the same module VL-to-VL:35.9 ± 7.7%, DM-to-DM: 29.4 ± 6.8%, mean ± SD) than between them, and ventrolateral module cells sending three times more projections to the contralateral dorsomedial cells than vice versa (VL-to-DM: 26.0 ± 5.4%, DM-to-VL: 8.5 ± 3.3%, two-way ANOVA with Tukey's post-hoc comparisons, letters indicate $p < 0.01$, $n = 7$ SCN). D, Dorsal; V, Ventral; M, Medial; L, Lateral; 3V, 3rd Ventricle; OC, Optic Chiasm.

#### Supplementary Figure 11



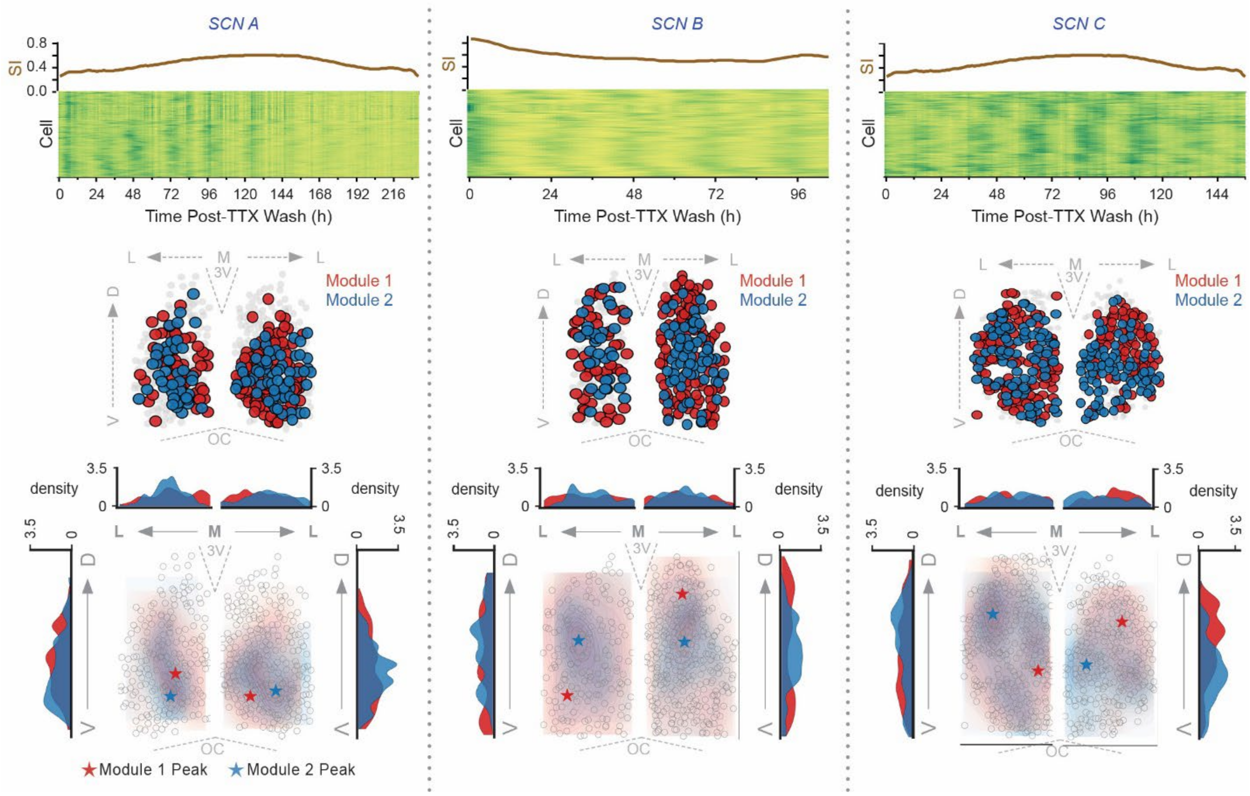


**Supplementary Fig. 11 | Dorsomedial and ventrolateral modules were not observed in SCN that failed to synchronize.** Raster plots and sync index (SI) from three SCN (SCN A-C) show how cellular PER2 rhythms failed to synchronize after tetrodotoxin (TTX) removal. Spatial maps of the three bilateral SCN (circles, middle) and density contours (bottom) show high inter-SCN variability in the two identified modules (red and blue), with many cells not assigned to any module (gray). Histograms show cell density along the dorsal-ventral (side) and medial-lateral SCN (top). D, Dorsal; V, Ventral; M, Medial; L, Lateral; 3V, third ventricle; OC, Optic Chiasm.

#### Supplementary Figure 12 **Nikhil et al**

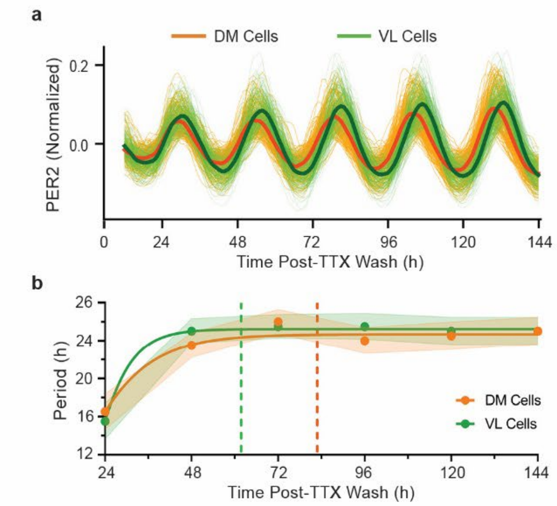


**Supplementary Fig. 12 | Ventrolateral SCN cells synchronize their circadian PER2 rhythms faster than the dorsomedial cells. a,** Normalized PER2 traces recorded from single SCN cells (thin lines) showed that, on average (thick lines), DM cells (orange) had higher daily amplitudes and peaked later than VL cells (green). **b,** Circadian period stabilized (< 0.5h deviation/day) earlier in VL cells (green, dashed line) compared to DM cells (orange, dashed line). Solid lines represent Gompertz curve fit; error bands are SD across cells.

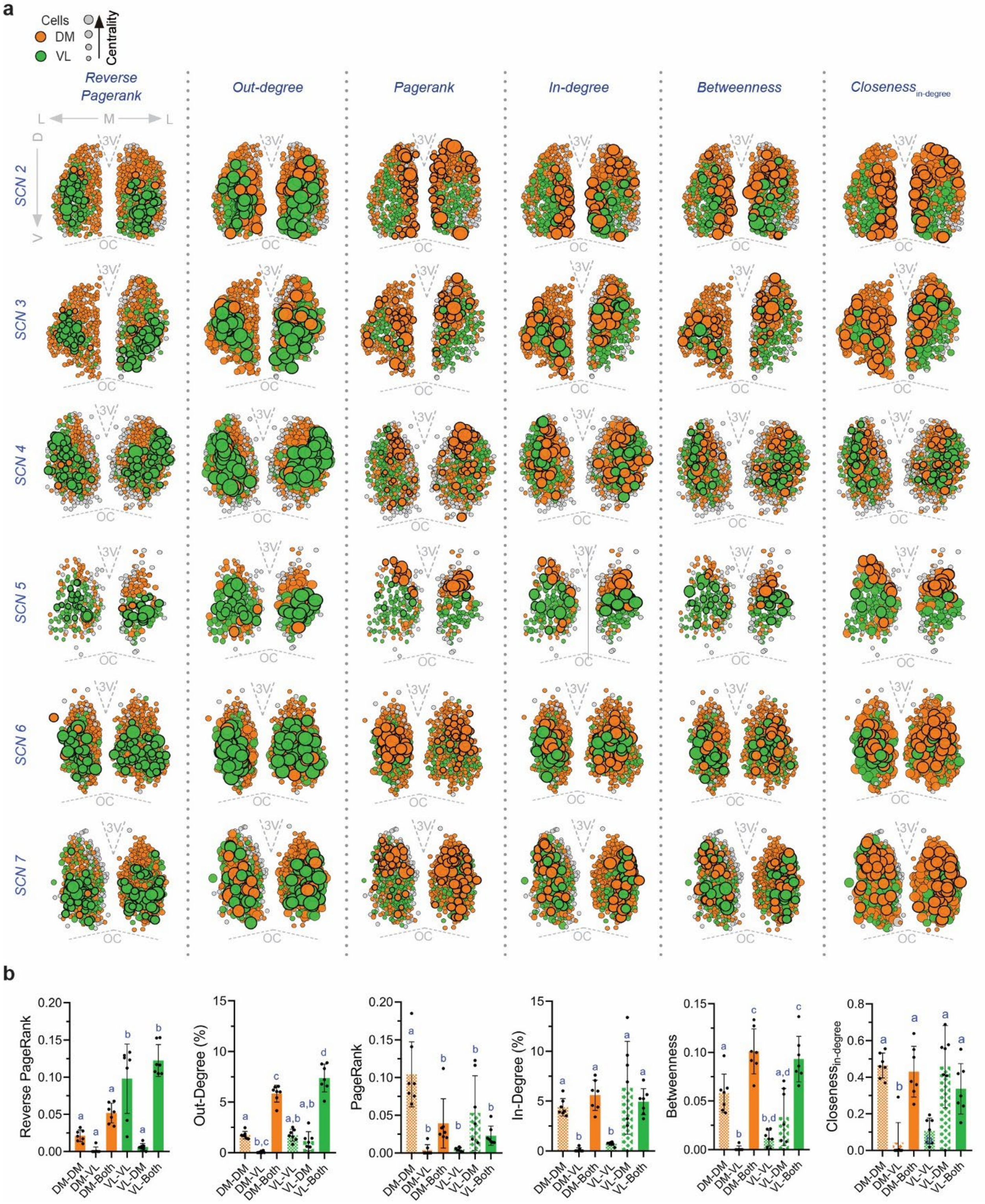


**Supplementary Fig. 13 | Ventrolateral cells are hubs with greater projections to other influential SCN cells.** Spatial maps of six other bilateral SCN show ventrolateral cells (VL, green circles) with high Reverse Pagerank (signal generators, larger cell = higher centrality, thick borders highlight the top 30% central cells)

compared to dorsomedial (DM, orange circles) cells. Similar representation of highly central cells based on Out-degree (broadcasters), In-degree (percent of the network from which a cell receives afferents), Betweenness (bridge cells mediating information flow), $\text{Closeness}_{\text{in-degree}}$ centralities (cells that are topologically closer and receive signals faster) and Pagerank centrality (sink cells to which signals ultimately flow). **b,** Mean Reverse Pagerank and other centralities of the dorsomedial and ventrolateral cells categorized based on their target modules (DM-DM: cells with dorsomedial to dorsomedial efferents, DM-VL: dorsomedial to ventrolateral: DM-Both: dorsomedial to both modules, VL-VL: ventrolateral to ventrolateral, VL-DM: ventrolateral to dorsomedial, VL-Both: ventrolateral to both modules; One-way ANOVA with Tukey's post hoc comparisons, Letters indicate $p < 0.01$, $n = 7$ SCN). Error bars are SD. D, Dorsal; V, Ventral; M, Medial; L, Lateral; 3V, third ventricle; OC, Optic Chiasm.

#### Supplementary Figure 14 **Nikhil et al**

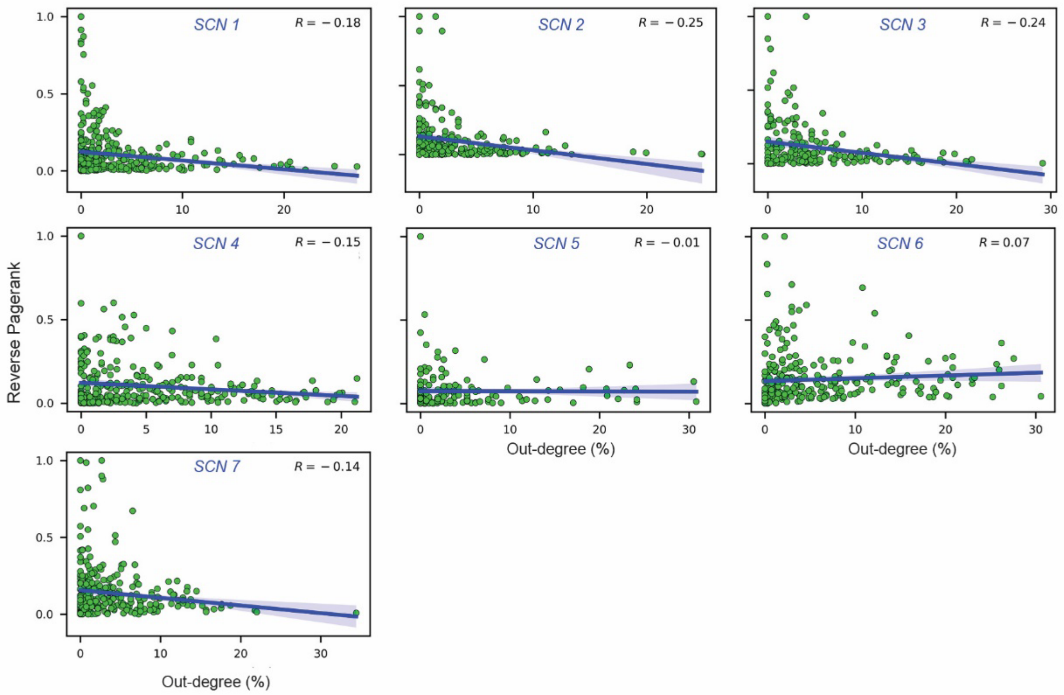


**Supplementary Fig. 14 | Signal generators and broadcasters in the ventrolateral SCN are distinct but with some overlap.** Linear fits (blue solid line, shaded region: 95% CI) show no correlation between Reverse Pagerank and Out-degree for VL SCN cells (green circles). *R*, Pearson correlation coefficient.

### Supplementary Figure 15 | Nikhil et al

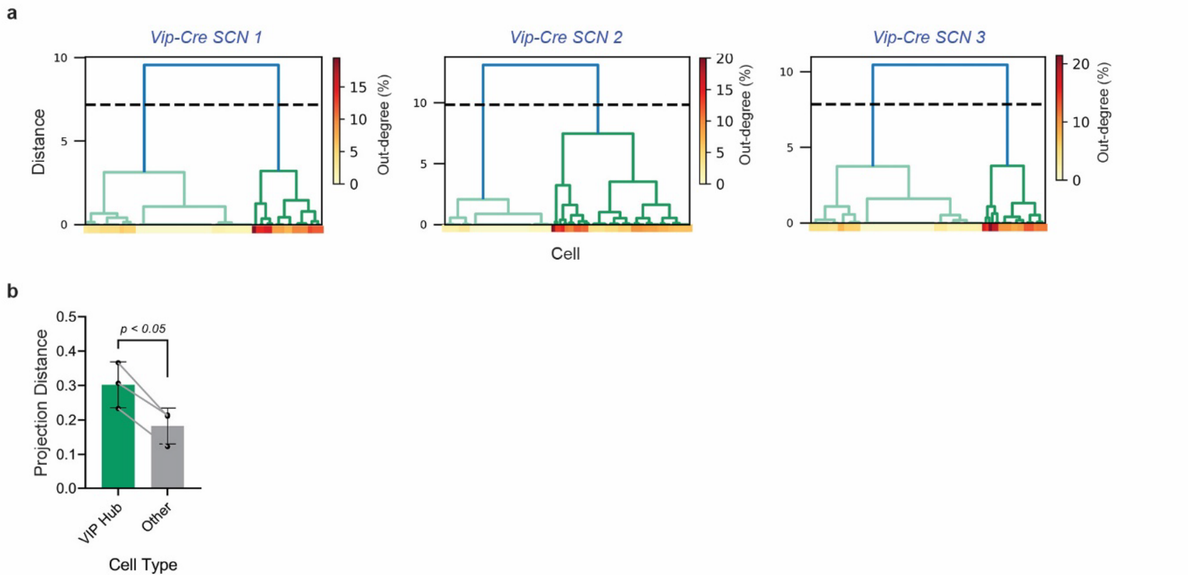


**Supplementary Fig. 15 | A subset of VIP neurons are hubs with higher numbers of efferents. a,** Unsupervised hierarchical clustering of VIP neurons by their out-degree (% of network a cell projects to) revealed two groups of VIP neurons consistently in three bilateral Vip-Cre SCN (SCN 1-3). One group of cells clustered with high out-degree (Hub neurons, dark green dendrogram, red saturation) and a second group of VIP neurons (non-Hub, light green dendrogram) with low out-degree (yellow saturation). The dendrogram was cut at 75% of maximal clustering distance (dashed line) to identify the two clusters. **b,** VIP Hub neurons had long-range functional projections (mean Euclidian distance of the cell to all its targets) compared to other SCN cells (paired Student's t-test, *n* = 3 bilateral SCN). Error bars are SD.

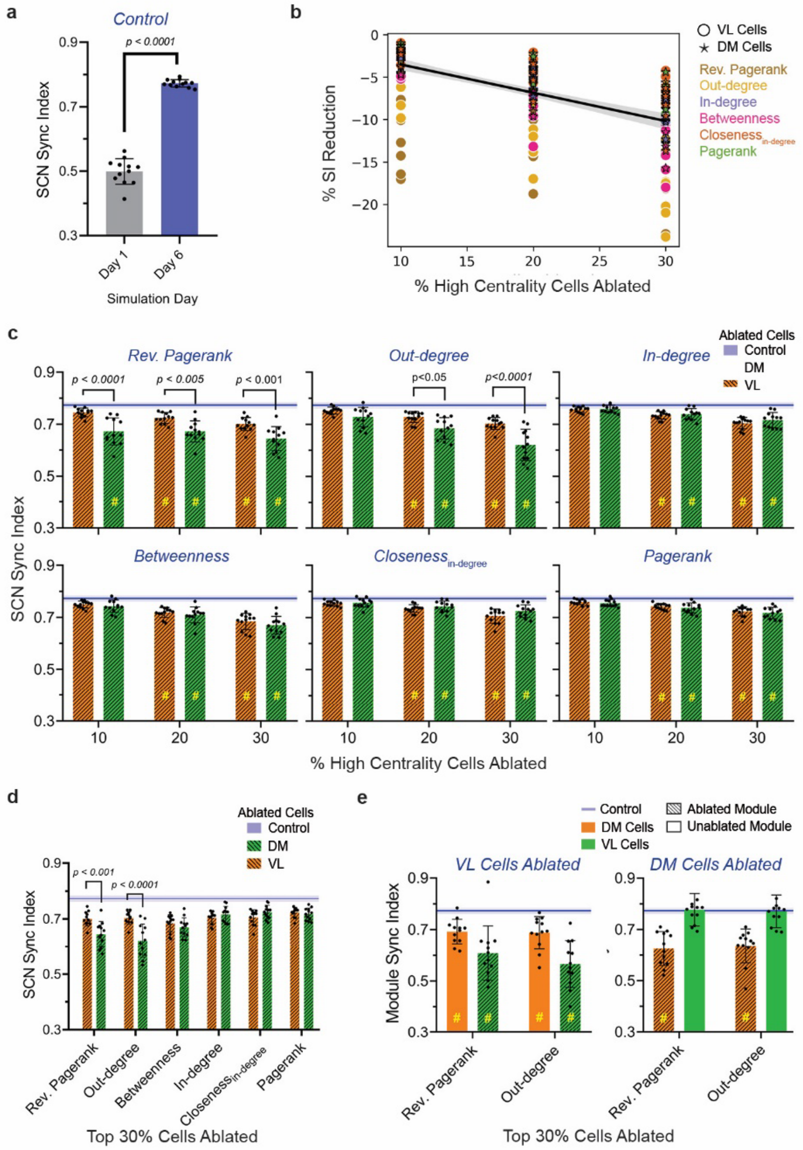


**Supplementary Fig. 16 | Ventrolateral cell ablation dose-dependently disrupts SCN synchronizability more than dorsomedial cells. a,** Synchrony index increased by day 6 of simulations illustrating the ability of cells to spontaneously synchronize from desynchronized conditions in 'control' SCN networks (no cells ablated; unpaired Student's t-test, using networks inferred from $n = 12$ unilateral SCN, one SCN could not be simulated within our parameter space). **b,** Average percentage reduction in sync index (relative to unablated controls calculated on day 6 following ablation of 10-30% of cells) modestly decreased even after 30% cell ablation ($R^2$: goodness of fit; $r$: Pearson correlation coefficient). **c,** Synchrony index (day 6 post-ablation) showed dose-dependent reduction in SCN synchronizability upon 10, 20, and 30% ablation of dorsomedial (DM, orange) and ventrolateral (VL, green) cells with different centralities: High Reverse Pagerank, Out-degree, Pagerank, In-degree, Betweenness, and Closeness centrality. **d,** Synchrony index on day 6 post-ablation of 30% ventrolateral cells showed higher impact of high Reverse Pagerank and Out-degree ventrolateral cells compared to dorsomedial cells. Synchrony Index of control (no cells ablated, blue line, error band = SD) in silico SCN (two-way ANOVA with Tukey's post-hoc comparisons, #: $p < 0.05$ for ablated vs. control comparisons, $n = 12$ unilateral SCN). **e,** Sync index of dorsomedial cells (orange) and ventrolateral cells (green) showed that ablating 30% high Reverse Pagerank or Out-degree dorsomedial cells (orange striped bars) reduced coherence only among dorsomedial cells but not the ventrolateral cells (green solid bars), whereas ablating ventrolateral cells (green striped bars) reduced synchrony among cells of both modules. Each data point is the average of 10 simulations. Error bars are SD.

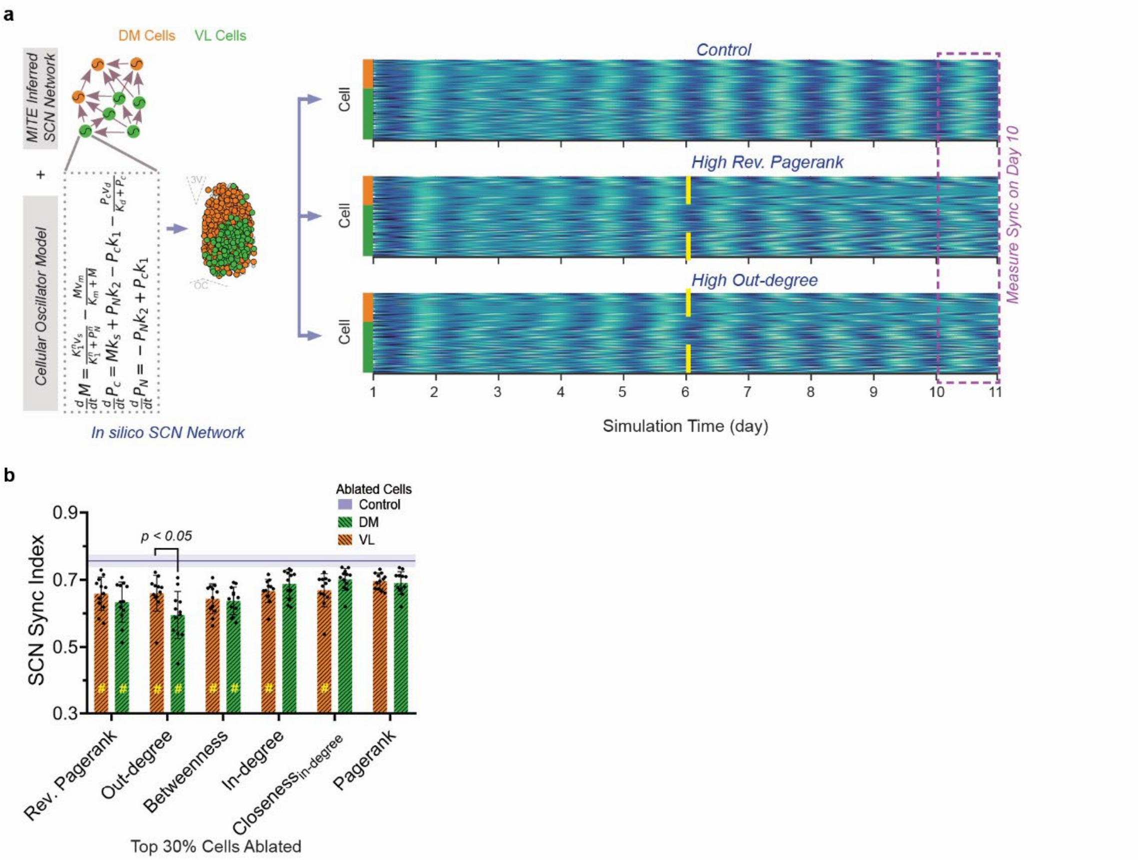


**Supplementary Fig. 17 | SCN cells that establish circadian synchrony also maintain synchrony. a,** In silico SCN networks were generated by coupling model cells with connection maps inferred by MITE from cellular PER2 recordings. Raster plots of PER2 expression from dorsomedial (DM, orange) and ventrolateral (VL, green) cells. In a representative simulation, cells in a control SCN showed increasing synchrony until day 6 which sustained until day 10 (top). In contrast, ablating 30% of ventrolateral cells with high Reverse Pagerank (middle) or Out-degree (bottom) on day 6 impaired sustained synchrony. **b,** SCN sync indices on day 10 of simulations showed how ablating VL cells with high Reverse Pagerank or Out-degree (orange striped bar) impaired the ability of the SCN to maintain circadian synchrony compared to control (blue line, error band = SD, two-way ANOVA with Tukey's post-hoc comparisons, #: $p < 0.05$ for ablated vs. control comparisons, $n$ = 12 unilateral SCN Error bars are SD. D, Dorsal; V, Ventral; M, Medial; L, Lateral; 3V, third ventricle; OC, Optic Chiasm.

**Supplementary Table 1 | Network statistics for unilateral and bilateral SCN**

| Metric | Nucleus | SCN 1 | SCN 2 | SCN 3 | SCN 4 | SCN 5 | SCN 6 | SCN 7 |
|---|---|---|---|---|---|---|---|---|
| # of Cells | Bilateral | 857 | 707 | 673 | 844 | 367 | 784 | 843 |
| | Left | 453 | 355 | 344 | 400 | 212 | 371 | 428 |
| | Right | 404 | 352 | 329 | 444 | 155 | 413 | 415 |
| Mean Degree | Bilateral | 3.06 | 3.41 | 3.81 | 3.92 | 3.35 | 4.08 | 3.53 |
| | Left | 3.09 | 3.28 | 3.68 | 3.93 | 3.14 | 4.33 | 3.53 |
| | Right | 2.95 | 3.61 | 3.90 | 4.01 | 4.03 | 3.87 | 3.55 |
| Path Length | Bilateral | 2.87 | 3.00 | 3.94 | 3.62 | 3.62 | 2.83 | 3.38 |
| | Left | 3.61 | 2.52 | 4.39 | 2.93 | 2.75 | 2.78 | 4.23 |
| | Right | 2.72 | 2.62 | 4.15 | 2.91 | 3.46 | 3.19 | 3.31 |
| # of Connections | Bilateral | 22424 | 16999 | 17212 | 27876 | 4495 | 25039 | 25066 |
| | Left | 6320 | 4116 | 4342 | 6280 | 1406 | 5939 | 6456 |
| | Right | 4806 | 4466 | 4213 | 7880 | 962 | 6591 | 6093 |
| # of Ipsilateral Connections | Bilateral | 13221 | 8922 | 9391 | 14262 | 3336 | 16780 | 14927 |
| # of Contralateral Connections | | 9203 | 8077 | 7821 | 13614 | 1159 | 8259 | 10139 |

**Supplementary Table 2 | Simulation parameters**

| Parameter | Description | Value |
|---|---|---|
| $K_1$ | mRNA transcription constant | 1 nM |
| $K_m$ | mRNA degradation constant | 0.5 nM |
| $n$ | mRNA transcription Hill term | 4 |
| $k_s$ | Protein translation rate | 0.417 1/h |
| $K_d$ | Cytosolic protein degradation constant | 0.13 nM |
| $v_d$ | Cytosolic protein degradation rate | 1.167 nM/h |
| $k_1$ | Cytosolic to nuclear protein rate | 0.417 nM/h |
| $k_2$ | Nuclear to cytosolic protein rate | 0.5 nM/h |
